# Targeting an anchored phosphatase-deacetylase unit restores renal ciliary homeostasis

**DOI:** 10.1101/2021.02.24.432764

**Authors:** Janani Gopalan, Mitch Omar, Ankita Roy, Nelly M. Cruz, Jerome Falcone, Katherine Forbush, Jonathan Himmelfarb, Benjamin S. Freedman, John D. Scott

**Affiliations:** Department of Pharmacology, University of Washington, Seattle, WA 98195, USA.; Kidney Research Institute, Division of Nephrology, Department of Medicine, University of Washington, Seattle, WA 98195, USA.; Institute for Stem Cell and Regenerative Medicine, and Department of Pathology, University of Washington, Seattle, WA 98195, USA

## Abstract

Pathophysiological defects in water homeostasis can lead to renal failure. Autosomal dominant polycystic kidney disease (ADPKD) is a common genetic disorder associated with abnormal cytoskeletal dynamics in the kidney collecting ducts and perturbed calcium and cAMP signaling in the ciliary compartment. We show that collecting ducts in mice lacking the A-Kinase anchoring protein AKAP220 exhibit enhanced development of primary cilia. Mechanistic studies reveal that AKAP220-associated protein phosphatase 1 (PP1) mediates this phenotype by promoting changes in the stability of histone deacetylase 6 (HDAC6) with concomitant defects in actin dynamics. This proceeds through a previously unrecognized adaptor function for PP1 as all ciliogenesis and cytoskeletal phenotypes are recapitulated in mIMCD3 knock-in cells expressing a phosphatase-targeting defective AKAP220-ΔPP1 mutant. Pharmacological blocking of local HDAC6 activity alters cilia development and reduces cystogenesis in kidney-on-chip and organoid models of ADPKD. These findings identify the AKAP220-PPI-HDAC6 pathway as a key effector in primary cilia development.

## Introduction

Kidneys recycle about 180 liters of fluid every day to partition nutrients and remove toxins from blood (Saborio et al., 2000). Water reabsorption from luminal fluid is triggered by the hormone arginine vasopressin via phosphorylation-dependent translocation of aquaporin-2 water pores to apical surface of kidney collecting ducts (Bankir et al., 2013; Noda et al., 2010; Yui et al., 2012). Not surprisingly, defects in renal water homeostasis have pathophysiological consequences. Approximately 35 million Americans suffer from chronic kidney diseases, characterized as a gradual loss of renal function (Hemmelgarn et al., 2006). Polycystic kidney diseases are disorders where the collecting ducts become enlarged with fluid filled cysts that reduce glomerular filtration rate (Wilson, 2004). Autosomal dominant polycystic kidney disease (ADPKD) with an estimated prevalence of 1 in 600 people, is a common genetic disorder associated with end-stage renal failure (Halvorson et al., 2010). Clinical evidence indicates that primary cilia function is altered in ADPKD (Ma et al., 2013, 2017). Hence this chronic renal disorder is classified as a ciliopathy (Badano et al., 2006; Fliegauf et al., 2007).

The primary cilium is a microtubule-based organelle protruding from the surface of most mammalian cells (Satir et al., 2010). In the kidney, primary cilia respond to fluctuations in fluid-flow through collecting ducts (Dell, 2015). They convert mechanical stimuli into biochemical signals to elicit developmental and regulatory responses (Mukhopadhyay et al., 2013; Somatilaka et al., 2020). Disease causing mutations in the ciliary transmembrane proteins polycystin 1 (*PKD1*) and polycystin 2 (*PKD2*) underlie ADPKD (Hughes et al., 1995; Mochizuki et al., 1996). Both proteins are components of a receptor-channel complex that responds to local second messenger signals (Harris and Torres, 2014). Accordingly, a ciliary hypothesis has been formulated that implicates defective calcium and cAMP signaling in the ciliary compartment as a factor in disease progression (Ma et al., 2017; Winyard and Jenkins, 2011).

A-kinase anchoring proteins (AKAPs) spatially constrain second messenger regulated kinases, protein phosphatases and GTPase effector proteins within subcellular compartments (Bucko and Scott, 2021; Langeberg and Scott, 2015; Omar and Scott, 2020; Smith et al., 2017, 2018; Taskén and Aandahl, 2004; Whiting et al., 2015). Several AKAPs participate in renal signaling, yet only a few anchoring proteins reside in cilia (Choi et al., 2011; Jo et al., 2001; May et al., 2020; Stefan et al., 2007). Ciliary AKAPs are postulated to position protein kinase A (PKA) as a negative regulator of hedgehog signaling (Breslow et al., 2018; Mukhopadhyay et al., 2013; Somatilaka et al., 2020). AKAP220 is a multifunctional anchoring protein that sequesters PKA, GSK3, the Rho GTPase effector IQGAP-1 and protein phosphatase 1 (PP1) (Logue et al., 2011; Schillace and Scott, 1999; Whiting et al., 2015). Each AKAP220-binding partner is implicated in local signaling events that potentiate ADPKD. PKA and PP1 bi-directionally control signaling in the ciliary compartment, whereas reduced Rho GTPase activity contributes to expansion of renal cysts (Parnell et al., 2012; Ye et al., 2017).

Here we show that AKAP220^-/-^ mice exhibit enhanced development of primary cilia. Mechanistic studies reveal that AKAP220 associated PP1 drives this phenotype by promoting changes in actin dynamics and histone deacetylase 6 (HDAC6) stability. Pharmacological blocking of local HDAC6 activity alters cilia development and reduces cystogenesis in cellular models of ADPKD. These findings point towards the AKAP220-PPI-HDAC6 pathway as a therapeutic target for the treatment of ADPKD.

## Results

### Loss of AKAP220 promotes renal cilia assembly

AKAP220 modulates cytoskeletal signaling events through its ability to recruit kinases, phosphatases and the small GTPase effector protein, IQGAP1 (Fig 1A; (Logue et al., 2011)). AKAP220^-/-^ mice display mild defects in water homeostasis and altered aquaporin 2 (AQP2) trafficking that are linked to disruption of an apical actin barrier in the kidney collecting ducts (Fig 1B; (Whiting et al., 2016)). Further investigation in kidney sections from wildtype and AKAP220^-/-^ mice led to the unexpected discovery that deletion of this anchoring protein correlated with increased numbers of primary cilia decorating each collecting duct (Fig 1C-H). The GTPase Arl13b (red) served as a ciliary marker, staining of AQP2 (green) marked kidney collecting ducts and DAPI (blue) highlighted nuclei (Fig 1C & D). Analysis of tissue sections collected from several animals are presented in Fig 1I, measuring a 3.5-fold increase in primary cilia in AKAP220^-/-^, as compared to wildtype. Additional measurements determined that cilia in AKAP220-/- collecting ducts are 1.68-fold longer than their wildtype counterparts (1.7 µm vs 1.02 µm; Fig 1J).

**Figure 1:**
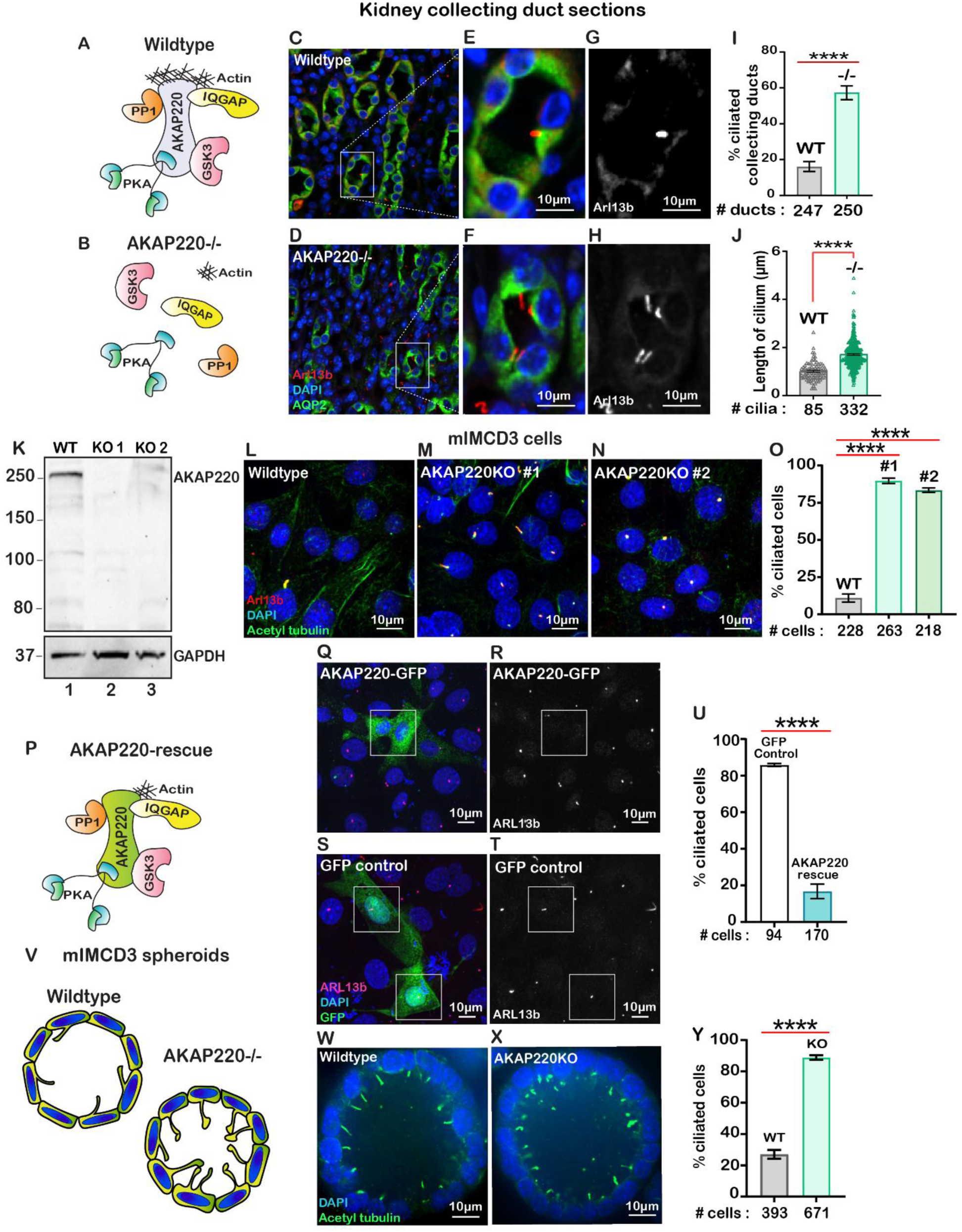
Loss of AKAP220 enhances ciliogenesis. **A**) Schematic of AKAP220 interaction with selected binding partners. **B**) Disruption of this signaling complex upon removal of AKAP220. Protein kinase A (blue), Glycogen synthase kinase-3 (pink), Protein phosphatase 1 (orange) and IQGAP (yellow) are indicated. **C-H**) Immunofluorescent staining of kidney collecting ducts with Arl13b (red), Aquaporin-2 (green) and DAPI (blue) from **C**) wildtype and **D**) AKAP220^-/-^ mice. **E & F**) Enlarged sections from wildtype and AKAP220^-/-^ mice. **G & H**) Grey scale images of Arl13b. **I**) Quantification (% ciliated collecting ducts) in wildtype (grey bar) and AKAP220^-/-^ (green bar). ****p<0.0001. **J**) Quantification of cilia length (µm) in wildtype (grey bar) and AKAP220^-/-^ (green bar). ****p<0.0001. Crispr-Cas9 gene editing of AKAP220 in mIMCD3 cells. **K**) Immunoblot detection of AKAP220 (top) and GAPDH loading control (bottom) from wildtype (lane 1) and AKAP220KO (lane 2) cell lysates. **L-N**) Immunofluorescent detection of primary cilia with acetyl tubulin (green), Arl13b (red) and DAPI (blue) in wildtype, and two independent clones of AKAP220KO mIMCD3 cells. **O**) Quantification (% ciliated cells) from wildtype (grey column), AKAP220KO#1 (green column) and AKAP220KO#2 (dark green column). ****p<0.0001, N=3. **P**) Schematic depicting reformation of the signaling complex upon rescue with AKAP220. Immunofluorescent detection of Arl13b (pink), GFP (green) and DAPI (blue) in **Q**) pEGFP-AKAP220 or **S**) GFP-control transfected AKAP220KO mIMCD3 cells. Grey scale image of Arl13b in **R**) control cells and **T**) AKAP220-rescued cells. **U**) Quantification (% ciliated cells) in pEGFP-AKAP220 (black bar) or GFP-control cells (teal bar). ****p<0.0001, N=3. **V**) Schematic of wildtype and AKAP220KO mIMCD3 spheroids. Immunofluorescent staining with acetyl tubulin (green) and DAPI (blue) in **W**) wildtype and **X**) AKAP220KO spheroids. **Y**) Quantification (% ciliated cells) in wildtype (grey column) and AKAP220KO (green column) spheroids. ****p<0.0001, N=3. All error bars are s.e.m. *P* values were calculated by unpaired two-tailed Student’s t-test. Scale bars (10µm). Number of cells analyzed indicated below each column.

Independent validation of this observation was provided when CRISPR-Cas9 gene-editing was used to delete AKAP220 from mouse Inner Medullary Collecting Duct (mIMCD3) cells. Immunoblot analysis confirmed the loss of AKAP220 in two independent clones (Fig 1K, top panel, lanes 2 & 3). GAPDH was used as loading control (Fig 1K, bottom panel). Immunofluorescent detection of primary cilia in each AKAP220KO clone measured an 8 to 10- fold increase in the percentage of ciliated cells in comparison to wildtype mIMCD3 cells (Fig 1L-O). Quantification is presented in Figure 1O. Serum starvation is typically used to ciliate mIMCD3 cells in culture (Westlake et al., 2011). However, serum starvation of AKAP220KO cell lines enhanced detection of multi-ciliated cells (supp Fig 1A-G). This made single cell quantification less accurate. For this reason, all further analyzes were conducted in asynchronous cultures.

Rescue experiments allowed us to attribute the increases in cilia to loss of AKAP220 (Fig 1P-U). AKAP220KO cells were transfected with a GFP-tagged AKAP220 construct (Fig 1P). Immunofluorescent detection of Arl13b (red) detected cilia and GFP-fluorescence confirmed rescue with the anchoring protein (Fig 1Q & R). GFP served as a control (Fig 1S & T). Quantification from three independent experiments confirmed that rescue with AKAP220 dramatically reduced the number of ciliated cells as compared to rescue with GFP (Fig 1U).

Since another anchoring protein AKAP150 has been detected in primary cilia, it was important to establish if this AKAP also contributed to primary cilia development (Choi et al., 2011). Gene-editing was used to generate mIMCD3 cells lacking AKAP150. Double knockout cells lacking AKAP150 and AKAP220 were also produced. Immunofluorescent detection of ARl13b (green) and acetyl-tubulin (red) as ciliary markers revealed that the loss of AKAP150 alone had no effect on cilia development (Supp Fig 1H & I). Yet, a pronounced increase in the ciliated population was observed in AKAP220/150 double knockout cells (Supp Fig 1J & K). These additional studies indicate that AKAP150 signaling does not support renal ciliogenesis.

Renal spheroids recapitulate features of ductal architecture for disease modelling ((Giles et al., 2014); Fig 1V). We cultured mIMCD3 cells in Matrigel for 72 hours to form three-dimensional spheroids (Fig 1W & X). Immunofluorescent detection of the ciliary marker acetyl-tubulin (green) confirmed that loss of AKAP220 correlated with persistence of primary cilia in mIMCD3 KO spheroids. Detection of DAPI (blue) marked nuclei (Fig 1W & X). Thus, local signaling mechanisms that proceed through AKAP220 impact ciliary development in cellular, three-dimensional spheroid and animal models of the renal microenvironment.

### AKAP220 contributes to histone deacetylase 6 stability

Primary cilia formation requires tubulin heterodimers that are stabilized by acetylation on lysine 40 (Portran et al., 2017). This prompted us to consider whether acetylated tubulin was abundant in cells lacking AKAP220. Experiments were conducted in two phases. First, immunofluorescent detection of acetylated alpha tubulin confirmed that this modified form was prevalent in AKAP220KO cells (Fig 2A & B). Second, immunoblot analysis showed that acetyl tubulin was more prominent in AKAP220KO cell lysates as compared to wildtype (Fig 2C, top panel). Alpha tubulin was consistent in both cell lines (Fig 2C, mid panel). GAPDH was the loading control (Fig 2C, lower panel). Amalgamated data from four independent experiments are quantified in figure 2D.

**Figure 2:**
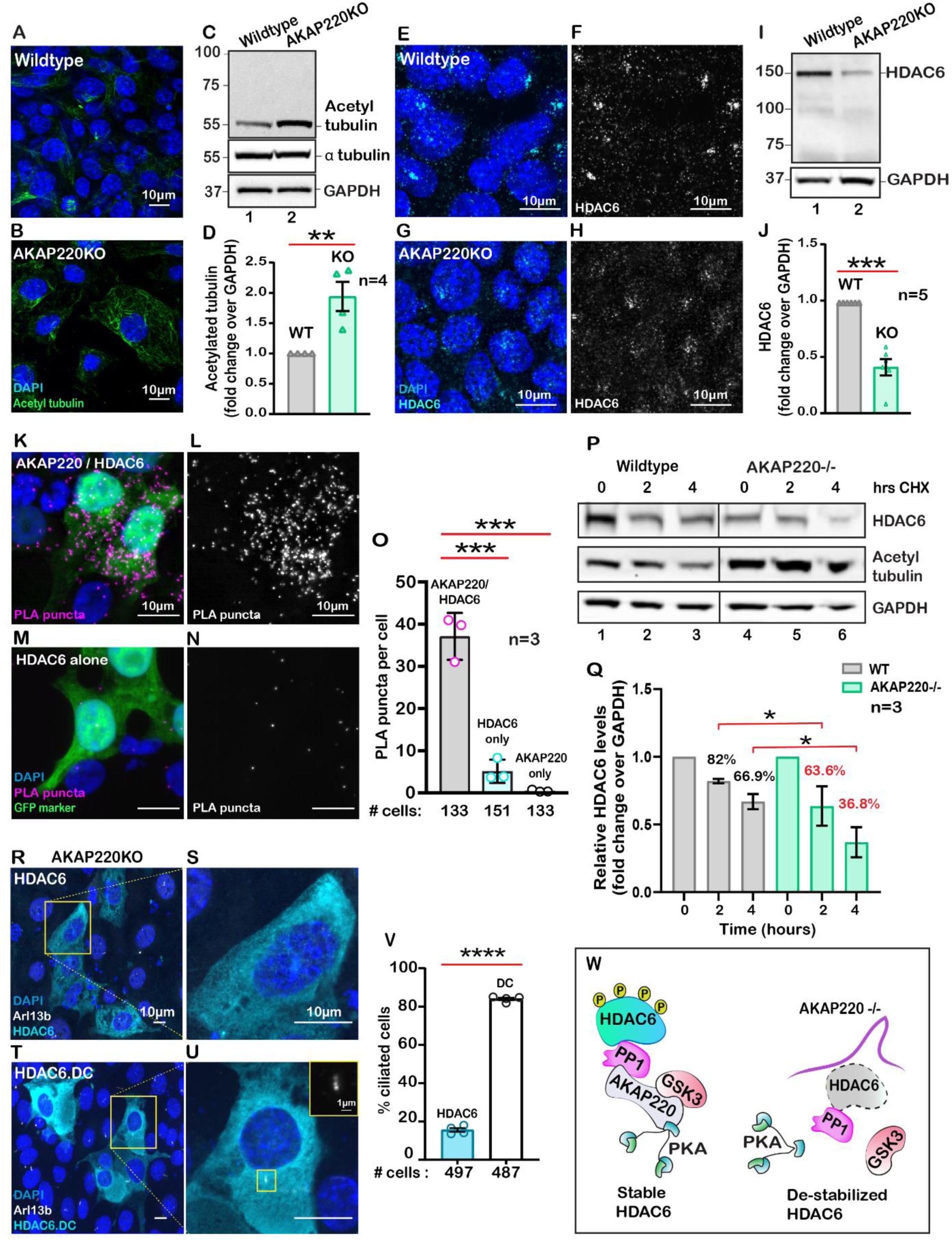
AKAP220 influences tubulin deacetylation. Immunofluorescent detection of acetyl tubulin (green) and DAPI (blue) in **A**) wildtype and **B**) AKAP220KO mIMCD3 cells. **C**) Immunoblot detection of acetylated tubulin (top), alpha tubulin (mid) and GAPDH loading control (bottom), in wildtype (lane 1) and AKAP220KO (lane 2) cell lysates. **D**) Quantification by densitometry of acetylated tubulin in wildtype (grey column) and AKAP220KO (green column) lysates. **p<0.01, N=4. Immunofluorescent staining of HDAC6 (cyan) and DAPI (blue) in **E**) wildtype and **G**) AKAP220KO cells. Grey scale images of HDAC6 in **F**) wildtype and **H**) AKAP220KO cells. **I**) Immunoblot detection of HDAC6 (top) and GAPDH as loading control (bottom) in wildtype (lane 1) and AKAP220KO (lane 2) cell lysates. **J**) Quantification by densitometry of HDAC6 in wildtype (grey column) and AKAP220KO (green column) lysates. ***p<0.001, N=5. **K**) Proximity ligation (PLA) detection of V5-AKAP220/HDAC6 subcomplexes (pink), DAPI (blue) in cells expressing GFP (green) as a transfection marker. **L**) Grey scale image highlights V5-AKAP220/HDAC6 PLA puncta. **M** & **N**) Control PLA experiments in cells treated with anti-HDAC6 antibody alone. **O**) Amalgamated data (PLA puncta/cell) from three independent experiments is presented. Cycloheximide pulse-chase assay investigated HDAC6 stability. **P**) Immunoblot of HDAC6 (top), acetylated tubulin (mid) and loading control GAPDH (bottom) from wildtype (lanes 1-3) and AKAP220KO (lanes 4-6) from mIMCD3 cells treated with cycloheximide. Data collected over a time course (0-4 h). **Q**) Quantification of amalgamated data by densitometry (N=3). HDAC6 levels in wildtype (grey) and AKAP220KO (green) are indicated. Levels of protein (%) are normalized to wildtype control (no treatment, 0hr). **R-V**) AKAP220KO mIMCD3 cells were transfected with **R**) flag-HDAC6 or **T**) catalytically inactive mutant HDAC6.DC. Immunofluorescent detection of HDAC6 (cyan), Arl13b (white) and DAPI (blue). Enlarged sections depict **S**) loss of primary cilium upon overexpression of active HDAC6 and **U)** intact cilium upon overexpression of inactive mutant HDAC6.DC. **V**) Quantification (% ciliated cells) in AKAP220KO cells transfected with flag-HDAC6 (black column) and HDAC6.DC (cyan column). ****p<0.0001, N=3. **W**) Schematic of how recruitment to the AKAP220-signaling complex stabilizes HDAC6 through local phosphorylation. All error bars are s.e.m. *P* values were calculated by unpaired two-tailed Student’s t-test. Scale bars (10µm). Number of cells analyzed indicated below each column.

Histone deacetylase 6 (HDAC6) catalyzes the de-acetylation of tubulin to promote depolymerization of primary cilia (Hubbert et al., 2002; Ran et al., 2015). Hence, we reasoned that the enhanced detection of acetylated tubulin in AKAP220KO cells may be a consequence of reduced HDAC6 activity. Five complementary methods tested this postulate. First, immunofluorescent detection of HDAC6 (cyan) confirmed its enhanced compartmentalization at the base of primary cilia in wildtype mIMCD3 cells (Fig 2E; Supp Fig 2A). Conversely, the subcellular distribution of HDAC6 was altered in AKAP220KO mIMCD3 cells (Fig 2G; Supp Fig 2C). This is most apparent in the grey scale images representing HDAC6 alone (Fig 2F & H, Supp Fig 2B & D). Second, immunoblot analysis of HDAC6 protein in mIMCD3 cell lysates detected reduced levels of the deacetylase in cells that lack AKAP220, as compared to wildtype controls (Fig 2I, top panel, lane 2). GAPDH served as a loading control (Fig 2I, bottom panel). Amalgamated data from five independent experiments indicated that deletion of AKAP220 correlated with a 50% reduction of total HDAC6 in mIMCD3 cells (Fig 2J). This argues that interface with AKAP220 serves to stabilize HDAC6.

Third, proximity ligation assay (PLA) detected AKAP220-HDAC6 complexes in situ. This approach identifies protein-protein interactions that occur within a range of 40–60 nm (Whiting et al., 2015). PLA puncta indicative of AKAP220/HDAC6 complexes were readily detected in cells transfected with V5-tagged AKAP220 (Fig 2K). Grey scale image emphasizes the distribution of AKAP220/HDAC6 complexes (Fig 2L). In contrast, detection of PLA puncta was dramatically reduced in control experiments performed in untransfected cells or in the absence of HDAC6 or V5 antibodies (Fig 2M & N, Supp Fig 2E-I). GFP was used as a transfection marker. Quantification from three independent experiments is presented (Fig 2O).

Fourth, we performed cycloheximide-chase experiments. Cells lysates were prepared at selected time points (0-4 hours; Fig 2P). Immunoblot detection of HDAC6 (top panel) and acetylated tubulin (mid panel) monitored the stability and activity of the de-acetylase over time. GAPDH served as a loading control (bottom panel). In wildtype mIMCD3 cells, HDAC6 protein was relatively constant and acetylated tubulin levels were low (Fig 2P, lanes 1-3). Conversely, upon loss of the anchoring protein, HDAC6 levels were markedly reduced over time and acetylated tubulin levels were elevated (Fig 2P, lanes 4-6). Densitometric analysis of HDAC6 levels in wildtype (grey) and AKAP220KO (green) from three independent experiments are presented (Fig 2Q). These results show a significant decrease in HDAC6 stability in the absence of the anchoring protein (Fig 2P & Q). Finally, overexpression of murine HDAC6 (cyan) abrogated primary cilia formation in AKAP220KO cells (Fig 2R & S). In contrast, expression of a catalytically inactive HDAC6 mutant (H216A, H611A) had no effect on cilia formation in the AKAP220 null background (Fig 2T & U). Amalgamated data from three experiments are presented (Fig 2V). Collectively, these studies implicate HDAC6 activity in AKAP220-mediated control of primary cilia development. Mechanistically, dephosphorylation of HDAC6 favors its degradation (Ran et al., 2020). Protein phosphatase 1(PP1) is a well-characterized binding partner of AKAP220 that also interacts with this deacetylase (Brush et al., 2004). Therefore, we reasoned that PP1 may serve as an adaptor protein that bridges HDAC6 to AKAP220 at the ciliary compartment (Fig 2W).

### AKAP220-anchored protein phosphatase 1 alters primary cilia turnover

Protein phosphatase 1 interacts with AKAP220 primarily through its KVQF motif (Schillace and Scott, 1999), and has recently been implicated in trafficking of polycystin-1 to cilia (Luo et al., 2019). We used CRISPR-Cas9 gene editing to generate a knock-in cell line where the phosphatase-targeting motif was replaced by a TATA sequence (Fig 3A & B; Supp Fig 3A & B). Co-immunoprecipitation assays confirmed that AKAP220-ΔPP1 is unable to anchor PP1 (Supp Fig 3C). Immunoblot analysis confirmed expression of AKAP220-ΔPP1 as compared to knockout control (Fig 3C, top panel, lane 2). GAPDH was a loading control (Fig 3C, bottom panel). To our surprise, characterization by immunofluorescence revealed that loss of a mere four-residue motif that enables PP1-anchoring to AKAP220 correlated with dramatically increased cilia numbers (Fig 3D & E). Arl13b (red) and acetyl tubulin (green) were used as ciliary markers and nuclei were detected by DAPI (blue). This striking phenotype is clearly portrayed in the gray scale images of Arl13b (Fig 3F & G). Additionally, three-dimensional surface plots show that primary cilia morphology is altered in the AKAP220-ΔPP1 cells, as compared to wildtype controls (Fig 3H & I). Results from at least three independent experiments are quantified (Fig 3N). Parallel analyses established that acetyl-tubulin was elevated in AKAP220-ΔPP1 (Fig 3J & L). This is highlighted in insets and 3D surface plots of acetylated tubulin (Fig 3K & M). Collectively, these results indicate that PP1 anchoring to AKAP220 is necessary for regulation of ciliary development and its loss is sufficient to drive enhanced ciliation. We reasoned that PP1 may serve as an adaptor protein that incorporates histone deacetylase 6 into AKAP220 signaling complexes.

**Figure 3:**
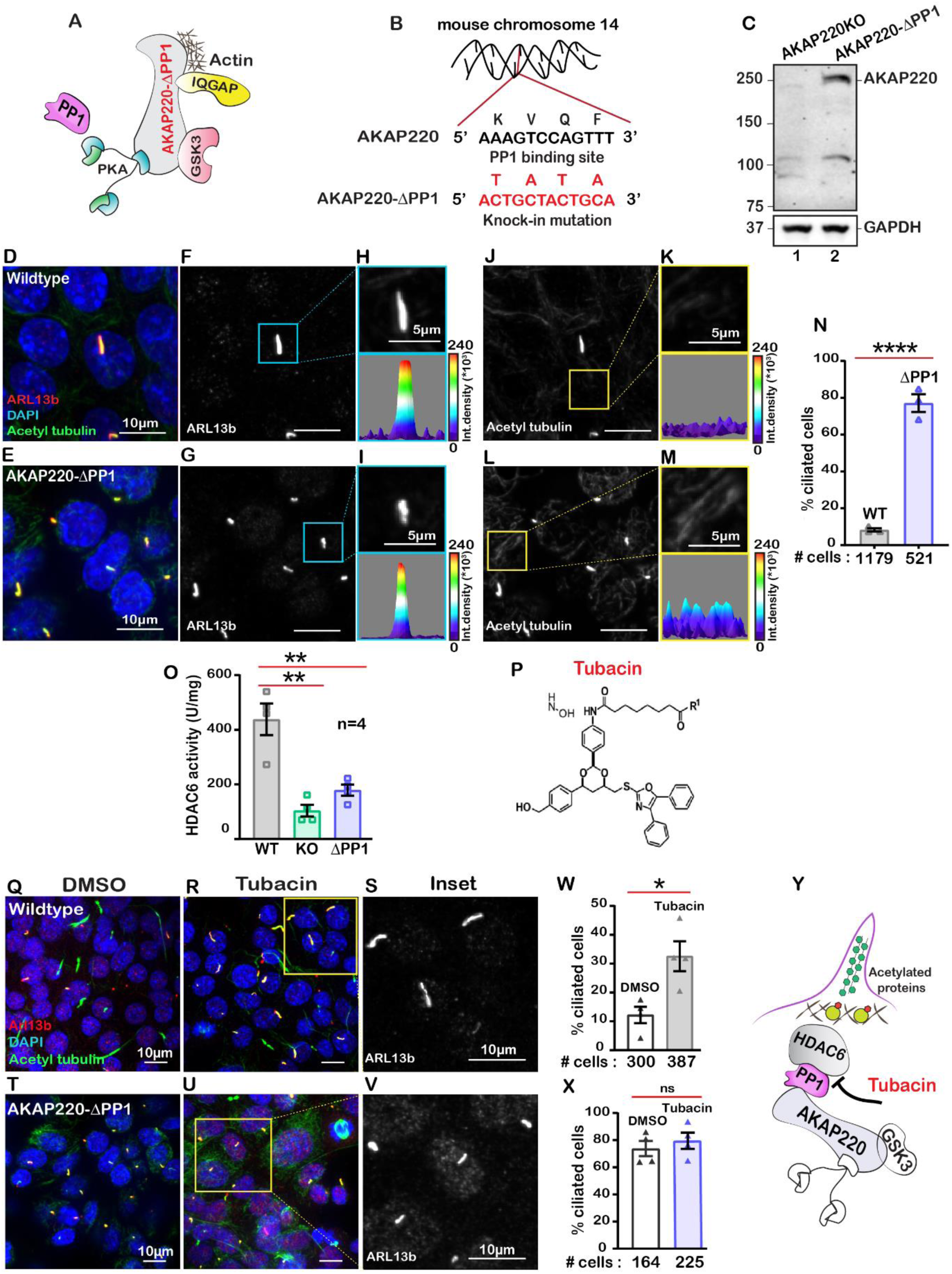
Anchored protein phosphatase 1 is necessary for HDAC6 activity. **A**) Schematic of AKAP220-ΔPP1. Binding partners are indicated. Gene editing deleted the principal phosphatase binding site (KVQF) on AKAP220. **B**) Nucleotide sequencing reveals substitution of the KVxF motif. **C**) Immunoblot detection of AKAP220 (top) and GAPDH loading control (bottom) in AKAP220KO (lane 1) and AKAP220-ΔPP1 (lane 2) mIMCD3 cell lysates. **D-N**) Immunofluorescent detection of acetyl tubulin (green), Arl13b (red) and DAPI (blue) in **D**) wildtype and **E**) AKAP220-ΔPP1 cells. Grey scale images of Arl13b in **F**) wildtype and **G**) AKAP220-ΔPP1 cells. A single enlarged cilium (top) and corresponding three-dimensional surface plots (bottom) from **H**) wildtype and **I**) AKAP220-ΔPP1 cells. Grey scale images of acetylated tubulin in **J**) wildtype and **L**) AKAP220-ΔPP1 cells. Enlarged sections from **J & L** (top) and corresponding three-dimensional surface plots (bottom) from **K**) wildtype and **M**) AKAP220- ΔPP1 cells. **N**) Quantification (% ciliated cells) in wildtype (grey) and AKAP220-ΔPP1 (blue). ****p<0.0001, N=3. **O**) HDAC6 activity levels (A.U.) in wildtype (grey), AKAP220KO (green) and AKAP220-ΔPP1 (blue) cells as assessed by Bioline’s activity assay. **p<0.01, N=4. **P**) Chemical structure of HDAC6 inhibitor tubacin. **Q-S**) Tubacin enhances ciliogenesis in the presence of native AKAP220. Wildtype mIMCD3 cells treated with **Q**) DMSO or **R**) tubacin (2µM) for 4 hrs. Immunofluorescent staining with acetyl tubulin (green), Arl13b (red) and DAPI (blue). **S**) Higher magnification grey scale image of Arl13b staining. **W**) Quantification (% ciliated cells) in DMSO (white) and tubacin-treated (grey) wildtype cells. *p<0.05, ns=non-significant; N=3. **T-V**) Tubacin has no effect on AKAP220-ΔPP1 cells. **T**) DMSO or **U**) and **V**) tubacin treated AKAP220-ΔPP1 cells. **X**) Quantification (% ciliated cells) and analysis as described above in DMSO (white) and tubacin-treated (blue). **Y**) Schematic of proposed tubacin mechanism of action on AKAP220- signaling complex. All error bars are s.e.m. *P* values were calculated by unpaired two-tailed Student’s t-test. Scale bars (10µm). Number of cells analyzed indicated below each column.

A fluorometric assay monitored HDAC6 activity in each cell type (Fig 3O). As expected, HDAC6 activity was reduced in AKAP220KO and AKAP220-ΔPP1, as compared to the wildtype mIMCD3 cells. Amalgamated data from four experiments are presented (Fig 3O). These findings suggest that the loss of HDAC6 activity and elevated acetylated tubulin correlate with persistence of primary cilia.

This led to a working hypothesis that AKAP220-targeted HDAC6 modulates primary cilia development. The HDAC6-selective inhibitor tubacin was used as a tool to evaluate the contribution of this enzyme activity in our mIMCD3 cell lines ((Haggarty et al., 2003); Fig 3P). This drug was applied to test if blocking anchored-HDAC6 activity enhanced ciliary development. In wildtype cells, application of tubacin (2µM) increased the number of ciliated cells as assessed by immunofluorescent detection of Arl13b (red) and acetylated tubulin (green; Fig 3Q & R). The grey scale image and quantification of three independent experiments reveal a 3.2-fold increase in percent ciliated cells upon pharmacologically targeting HDAC6 (Fig 3S & V). Co-staining with both cilia markers was necessary to delineate between cilia and acetyl-tubulin at the midbodies of dividing cells (Fig 3Q). Importantly, no change in cilia number occurred when AKAP220-ΔPP1 cells were treated with tubacin (Fig 3T-V). Similarly, AKAP220KO cells were unresponsive to the drug (Supp Fig 3H-K). This raises the intriguing possibility that an HDAC6-PP1-AKAP220 signaling axis is the intracellular target for the drug tubacin (Fig 3Y).

### AKAP220 signaling impacts cortical actin dynamics

The cytoskeleton maintains cell shape and structure by synchronizing the assembly and disassembly of actin, intermediate and tubulin filaments (Janke and Magiera, 2020; Klymkowsky, 1999). Covalent modification of these elements is an important facet of cytoskeletal regulation (Portran et al., 2017). For example HDAC6 deacetylates the actin-binding protein cortactin to relocate it from the nucleus to its sites of action (Ran et al., 2015). Since HDAC6 activity is impaired in AKAP220KO and AKAP220-ΔPP1 cells, it was important to evaluate if nuclear accumulation of acetylated cortactin was enhanced. Immunofluorescent detection of acetyl- cortactin (green) was more prominent in the nuclei of AKAP220KO and AKAP220-ΔPP1 mIMCD3 cells, as compared to wildtype (Fig 4A, C & E). This is emphasized in insets and corresponding surface plots (Fig 4B, D & F). Amalgamated data from three experiments are presented (Fig 4G). Next, we investigated if AKAP220 signaling influences actin filament morphology (Fig 4H-M). Immunofluorescent analyses show that AKAP220KO and AKAP220-ΔPP1 cells exhibit dramatic changes in the actin cytoskeleton (red) (Fig 4H, J & L). Line plot analyses (40-50 cells) reveal that total loss of the anchoring protein or disruption of HDAC6-PP1 attachment correlates with enhanced accumulation of cortical actin (Fig 4I, K & M). These findings pointed towards defects in actin dynamics. Immunofluorescence data depicting actin distribution in confluent cultures of wildtype and AKAP220KO mIMCD3 cell lines is shown in supplemental figure 4A-D.

**Figure 4:**
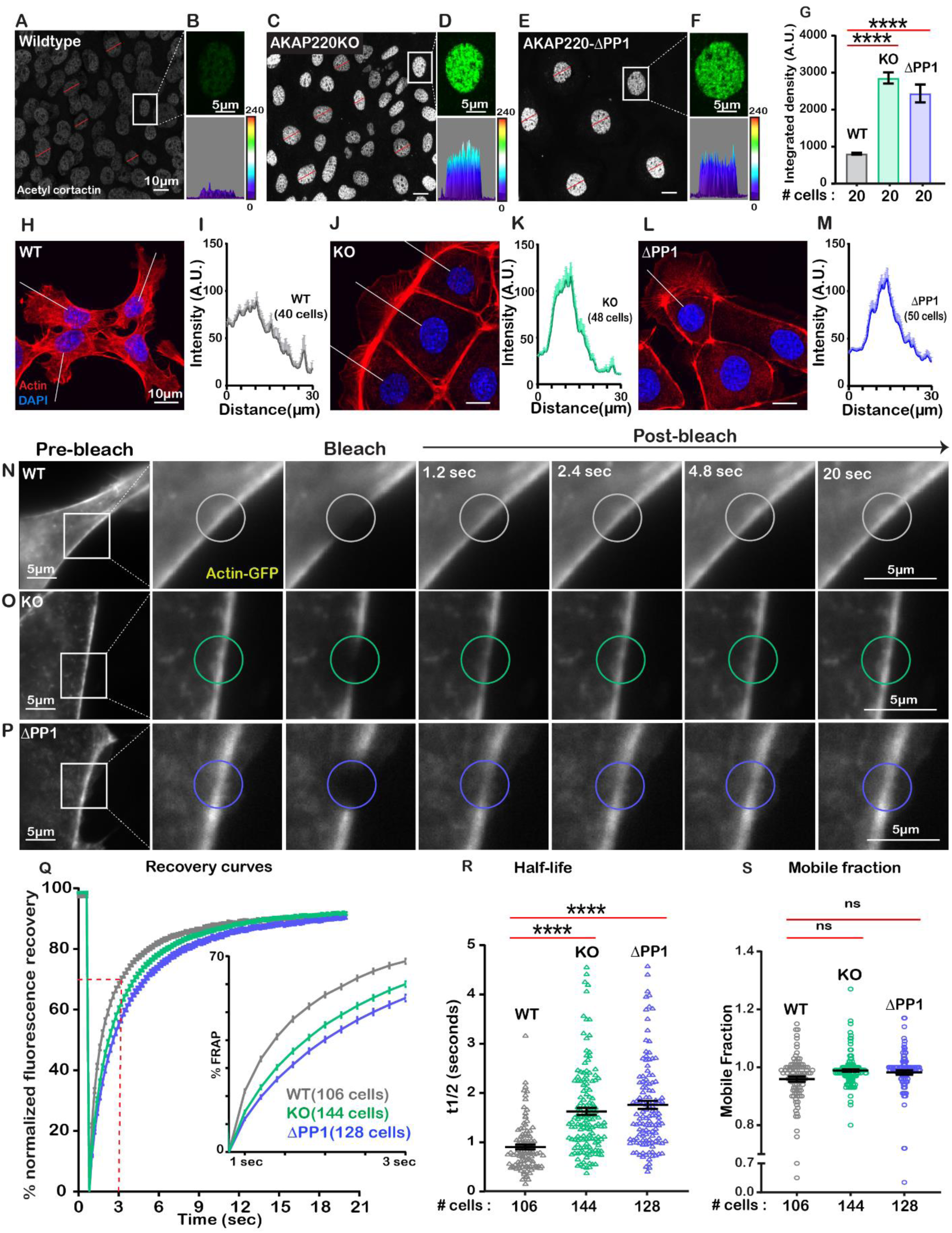
Loss of AKAP220-PP1 subcomplex impacts actin reorganization. Grey scale images of acetylated cortactin in **A**) wildtype, **C**) AKAP220KO and **E**) AKAP220-ΔPP1 mIMCD3 cells. Magnified images of nuclei (top) and three-dimensional surface plots (bottom) of acetylated cortactin in **B**) wildtype, **D**) AKAP220KO and **F**) AKAP220-ΔPP1 cells. **G**) Quantification of amalgamated data (20 cells) from wildtype (grey), AKAP220KO (green) and AKAP220-ΔPP1 (blue) cells. ****p<0.0001, N=3. Scale bars (10µm). **H-M**) Confocal images of actin (red) and DAPI (blue) in **H**) wildtype, **J**) AKAP220KO and **L**) AKAP220-ΔPP1 cells. Lines indicate sites of line plot analysis to measure actin distribution from nuclei to the lamellipodia in **I**) wildtype (grey), **K**) AKAP220KO (green) and **M**) AKAP220-ΔPP1 (blue) cells. Scale bars (10µm). **N-S**) Fluorescence recovery after photobleaching (FRAP) in mIMCD3 cells. Time course (0-20 sec) of GFP-actin imaging in **N**) wildtype, **O**) AKAP220KO and **P**) AKAP220-ΔPP1 cells. **Expanded section**) Photobleached portion of cortical actin. **Q**) FRAP curves in wildtype (grey), AKAP220KO (green) and AKAP220-ΔPP1(blue) cells. **Inset**) Photo recovery rates over the first 3 seconds. **R**) The t1/2 value for each cell analyzed is presented for wildtype (grey), AKAP220KO (green) and AKAP220-ΔPP1(blue) cells. ****p<0.0001, N=3. **S**) The mobile fraction of each cell analyzed is presented for wildtype (grey), AKAP220KO (green) and AKAP220-ΔPP1(blue circles) cells. Non-significant, N=3. Scale bars (5µm). All error bars are s.e.m. *P* values were calculated by unpaired two-tailed Student’s t-test. Number of cells analyzed indicated below each column.

To further explore this concept, we performed <u>F</u>luorescence <u>R</u>ecovery <u>a</u>fter <u>P</u>hotobleaching (FRAP) at cell-cell junctional actin fibers (Mov 1-3). Actin-GFP recovery upon photobleaching was monitored over a time course of 20 seconds in wildtype (grey; Fig 4N); AKAP220KO (green; Fig 4O) and AKAP220-ΔPP1(blue; Fig 4P) mIMCD3 cells. Recovery curves depict the half-life and mobile fraction (Fig 4Q). Expanded section accentuates the expedited recovery of actin in wildtype cells (Fig 4Q, inset). The t1/2 for photo-recovery is 1.6 sec; n=106 (Fig 4R, grey). In contrast, the t1/2 was 0.9 sec; n=144 for AKAP220KO cells (Fig 4R, green). This is a 0.72 sec ±0.09 reduction in the rate of photo-recovery when compared to wildtype. Parallel FRAP experiments in AKAP220-ΔPP1 cells calculated a t1/2 of 1.75 sec; n=128, representing a 0.85 ± 0.10 sec reduction in the rate of photo-recovery as compared to wildtype (Fig 4R, blue). The availability of actin-GFP was similar in mobile fractions from each cell type (wildtype 95% (grey); AKAP220KO 98% (green); AKAP220-ΔPP1 98% (blue); Figure 4S). Thus, disruption of AKAP220-signaling impacts the distribution and dynamics of actin filaments.

### Actin polymerization dictates cilium biogenesis and length

Actin is a key regulator of cilia formation and elongation (Kim et al., 2015). AKAP220^-/-^ mice exhibit reduced accumulation of apical actin through the diminished GTP-loading of RhoA (Whiting et al., 2016). This key regulator of cytoskeletal reorganization contributes to the assembly of actin barriers in renal cells (Blattner et al., 2013). HDAC6-mediated stimulation of actin polymerization is a prerequisite for the disassembly of primary cilia (Ran et al., 2015). A convergence of these ideas hypothesizes that defective AKAP220-signaling alters the dynamics of the actin barrier assembly to enhance ciliation (Fig 5A).

**Figure 5:**
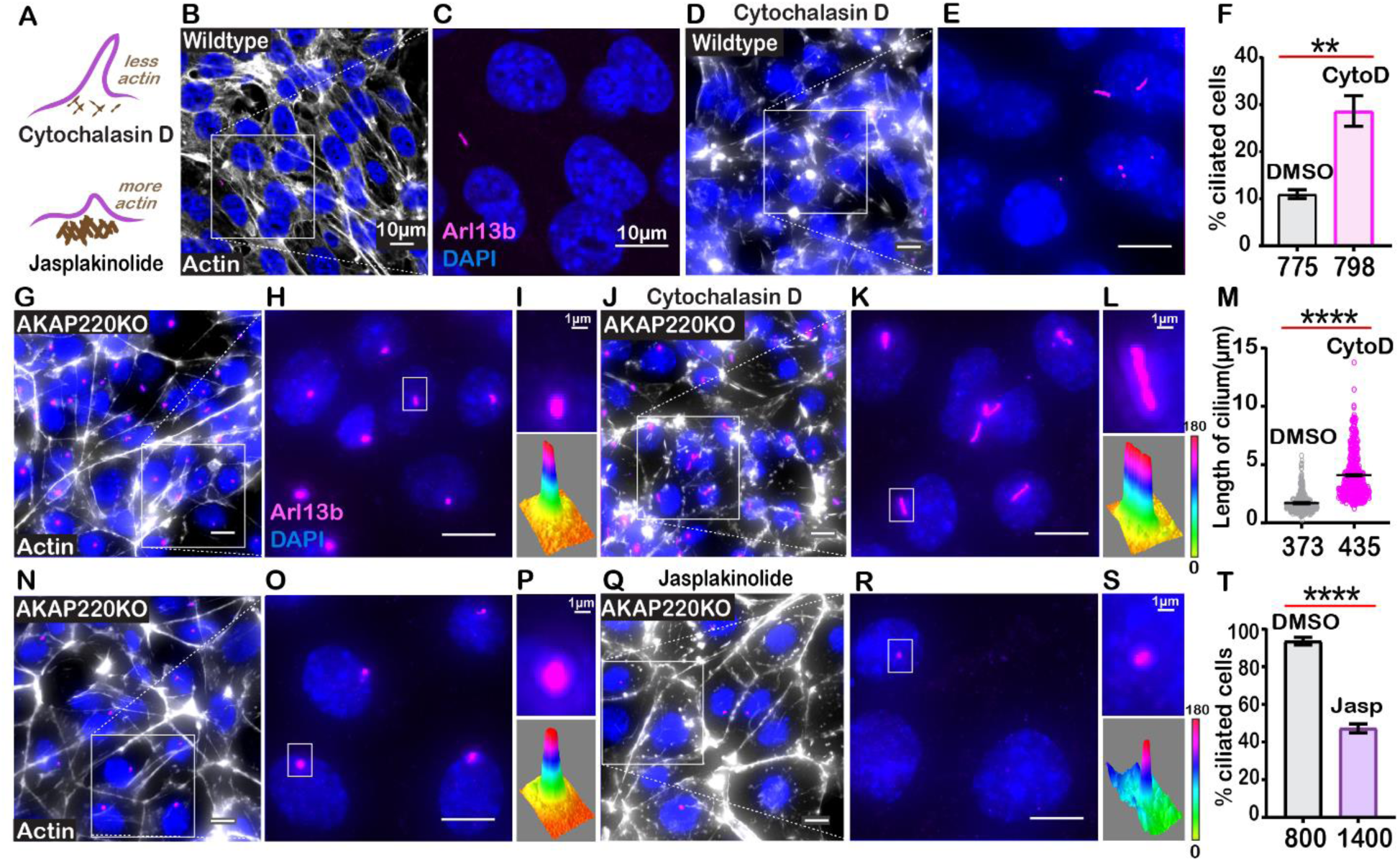
Cilia frequency and development are f-actin dependent. **A**) Schematic of how actin modulating drugs impact primary cilia. Cytochalasin D depolymerizes actin barriers. Jasplakinolide stabilizes actin filaments. **B-F**) Immunofluorescent detection of actin (white), Arl13b (pink) and DAPI (blue) in wildtype mIMCD3 cells treated with **B**) DMSO or **D**) 200nM Cytochalasin D. Enlarged regions emphasize cilia frequency in **C**) DMSO and **E**) Cytochalasin D-treated cells. **F**) Quantification (% ciliated cells) in DMSO (grey) and Cytochalasin D (pink) treated cells. **p<0.01, N=3. **G-M**) Immunofluorescent detection of actin (white), Arl13b (pink) and DAPI (blue) in **G**) DMSO and **J**) Cytochalasin D treated AKAP220KO cells. **Inset)** Expanded field of cells treated with **H**) DMSO or **K**) Cytochalasin D. Boxed regions in **I** and **L** focus on a single cilium (top) and three-dimensional surface plot (bottom). The width of the cylindrical region in the 3D surface plot represents cilium length. **M**) Quantification of cilia length in DMSO (grey) and Cytochalasin D (pink) treated cells. ****p<0.0001, N=3. **N-T**) Immunofluorescent staining of actin (white) and DAPI (blue) of **N** (DMSO) and **Q** (Jasplakinolide) treated AKAP220KO cells. **Inset**) Expanded field of cells treated with **O**) DMSO or **R**) Jasplakinolide. Boxed regions in **P** and **S** focus on a single cilium (top) and three-dimensional surface plot (bottom). The width of the cylindrical region in the 3D surface plot represents cilium length. **T**) Quantification (% ciliated cells) in DMSO (grey) and Jasplakinolide (purple). ****p<0.0001, N=3. All error bars are s.e.m. *P* values were calculated by unpaired two-tailed Student’s t-test. Scale bars (10µm). Number of cells analyzed indicated below each column.

To test this hypothesis, we treated wildtype cells with the actin-depolymerizing drug Cytochalasin D (Fig 5B-E). Drug application (200 nM) for 4 h favored disassembly of actin (white) as monitored by immunofluorescence (Fig 5B & D). Cytochalasin D promoted a 2.95-fold increase in the percentage of ciliated cells (Fig 5F, pink column). Similar effects were observed in AKAP220KO cells (Fig 5G-L). These pharmacological effects were scored by measuring the length of cilia (Fig 5M, pink). Insets feature representative cilia at higher magnification (Fig 5I & L, upper panels). Three-dimensional surface plots depict drug-induced changes in cilia length (Fig 5I &L, lower panels). Thus, pharmacological blockade of actin assembly augments ciliogenesis. This effect is enhanced in AKAP220 null cells.

Jasplakinolide is a macrocyclic peptide that stabilizes f-actin and augments the formation of actin barriers (Holzinger, 2009). Drug treatment (500 nM) for 1.5 h enhanced detection of cortical actin (white) and decreased the number of ciliated cells (pink; Fig 5N-S). Insets feature representative cilia at higher magnification (Fig 5P & S, upper panels). Three-dimensional surface plots depict drug-induced changes in cilia length (Fig 5P & S, lower panels). Collectively, these pharmacological studies show that bi-directional modulation of actin dynamics directly affects ciliation. This suggests that the actin dynamics changes observed in AKAP220KO and AKAP220- ΔPP1 cells underlie enhanced ciliation.

### AKAP220 signaling influences cilia morphology

We reasoned that a consequence of altered actin dynamics could be changes in cilia morphology. Super-resolution immunofluorescence imaging of fixed cells was performed using Arl13b as a ciliary marker. Cilia appeared retracted in AKAP220-null and ΔPP1-knock in cells in comparison to cylindrical and symmetrical architecture of cilia in wildtype cells (Fig 6A-D). A secondary feature was bulbous tips at the distal end of mutant cilia (Fig 6C & D). Analyses from three independent experiments show that the occurrence of bulbous tips was prevalent in cilia from AKAP220KO and AKAP220-ΔPP1 cells (Fig 6E).

**Figure 6:**
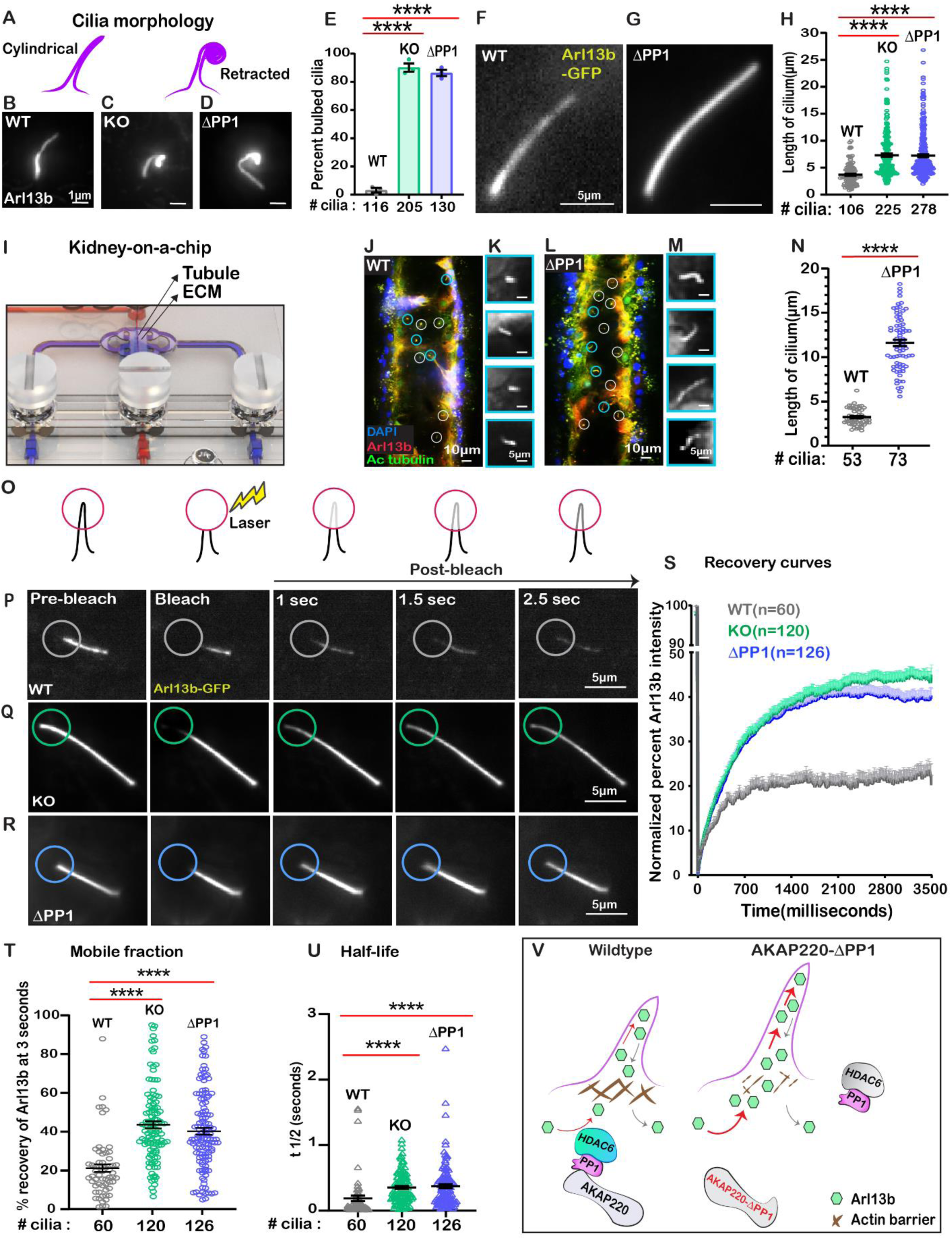
Loss of phosphatase anchoring promotes cilium elongation. **A**) Schematic of cylindrical and retracted cilia morphologies (purple). Super-resolution fixed-cell imaging of Arl13b (grey scale) in **B**) wildtype, **C**) AKAP220KO and **D**) AKAP220-ΔPP1 cilia. **E**) Quantification (% bulbed cilia) in wildtype (grey), AKAP220KO (green) and AKAP220-ΔPP1 (blue) cells. ****p<0.0001, N=3. Scale bars (1µm). Super-resolution live-cell images of Arl13b-GFP in **F**) wildtype and **G**) AKAP220-ΔPP1 cilia. **H**) Quantification of cilia length in wildtype (grey), AKAP220KO (green) and AKAP220-ΔPP1 (blue) cells. ****p<0.0001, N=3. Scale bars (5µm). **I**) Kidney-on-a-chip device. Location of kidney tubule (Tubule) and extracellular matrix (ECM) are indicated. Confocal imaging of **J**) Wildtype and **L**) AKAP220-ΔPP1cells cultured in chip device with Arl13b (red), acetyl tubulin (green) and DAPI (blue). Cilia (white circles) are marked. **Insets**) Magnified images of representative cilia (cyan circles) from **K**) wildtype and **M**) AKAP220-ΔPP1 pseudotubules. **N**) Quantification of cilia length in wildtype (grey) and AKAP220-ΔPP1 (blue). ****p<0.0001, N=3. Scale bars (10µm). Inset scale bars (5µm). **O-U**) Fluorescence recovery after photobleaching (FRAP) of Arl13b-GFP in primary cilia. **O**) Schematic of how FRAP was measured. **P-R**) Super-resolution live-cell images of Arl13b-GFP in **P**) wildtype, **Q**) AKAP220KO and **R**) AKAP220-ΔPP1 cells. Circles mark bleached portion of cilia. **S**) FRAP recovery curves of Arl13b over time (3500 ms) in wildtype (grey), AKAP220KO (green) and AKAP220-ΔPP1 (blue) cells. Scale bars (5µm). **T**) Quantification of mobile fraction in wildtype (grey), AKAP220KO (green) and AKAP220-ΔPP1 (blue) cilia. ****p<0.0001, N=3. **U**) Half-life of Arl13b in wildtype (grey), AKAP220KO (green) and AKAP220-ΔPP1 (blue) cilia. ****p<0.0001, N=3. **V**) Schematic depicting how AKAP220 modulation of the actin barrier influences movement of proteins in and out of primary cilia. All error bars are s.e.m. *P* values were calculated by unpaired two-tailed Student’s t-test. Number of cilia analyzed indicated below each column.

To investigate this phenomenon further in living cells, we infected our mIMCD3 cell lines with a lentiviral vector encoding Arl13b-GFP (Mov 4-6; Fig 6F-H). Quantitative imaging by live-cell super-resolution microscopy revealed that cilia in AKAP220KO (green) and AKAP220-ΔPP1 (blue) cells were 1.6 and 1.7-fold longer than wildtype counterparts (grey; Fig 6H). This led us to the conclusion that the bulbous tips presented in figures 6C & D were elongated, flexible cilia that partially retracted (coiled back) on themselves (Supp Fig 6).

Kidney-on-a-chip technology offers a pseudo-physiological environment that simulates kidney tubules (Weber et al., 2016). Microfluidic delivery of nutrients through the lumen of these tubules recapitulates fluid-flow (Freedman et al., 2013). This sophisticated tissue-engineering approach was used to evaluate cilia development (Fig 6I). Culturing of wildtype mIMCD3 cells formed columnar organoids with few cilia protruding into the lumen (Fig 6J & K). In contrast, longer cilia were evident in AKAP220-ΔPP1 organoids (Fig 6L & M). Immunofluorescent staining of Arl13b (red) and acetyl tubulin (green) marked cilia and DAPI (blue) detected nuclei. The average cilia length was 3.95-fold greater in AKAP220-ΔPP1 pseudo-tubules (n=73 cilia) as compared to wildtype (n=53 cilia). These effects are more visible in the grey scale images of Arl13b alone (Fig 6K&M). Amalgamated data from three independent experiments are presented in figure 6N.

Cilia assembly requires the passage of materials through an actin barrier formed across the basal body (Farina et al., 2016). We reasoned that the dynamics of this process may be altered upon manipulation of AKAP220 signaling. Therefore, we combined super-resolution microscopy with <u>F</u>luorescence <u>R</u>ecovery <u>a</u>fter <u>P</u>hotobleaching (FRAP) to visualize GFP-Arl13b trafficking into individual cilia (Fig 6O). Arl13b recovery upon photobleaching was monitored over a time course of 2.5 seconds in wildtype (grey; Fig 6P); AKAP220KO (green; Fig 6Q) and AKAP220-ΔPP1(blue; Fig 6R) mIMCD3 cells. In wildtype cells, the rate of GFP-Arl13b recovery was steady over this time-course. FRAP was more rapid and robust in AKAP220KO and AKAP220-ΔPP1 cilia (Fig 6S). Total Arl13b recovered in AKAP220KO is increased 22.35 ±3% (n= 120 cilia; green; Fig 6T) over wildtype (n=60 cilia; grey; Fig 6T). Similarly, recovery of Arl13b in AKAP220-ΔPP1 cilia is 19.04 ± 3% increased (n=126 cilia; blue; Fig 6T) over wildtype. The t1/2 of Arl13b recovery was 178.9 milliseconds in wildtype cilia (grey), as compared to 344.7 and 361.6 milliseconds in AKAP220KO and AKAP220-ΔPP1 cilia respectively (Fig 6U; green and blue). These studies suggest that there is a larger mobile pool of Arl13b trafficking into the mutant cilia. One plausible mechanism is that cilia in AKAP220KO and AKAP220-ΔPP1 cells lack an architectural checkpoint that gates protein movement into this organelle. A likely candidate is an f-actin barrier at the base of the cilium (Fig 6V).

### HDAC6 inhibition attenuates renal cyst formation

Autosomal dominant polycystic kidney disease (ADPKD) is associated with mutations in *PKD1* and *PKD2,* changes in apical actin and cilia dysfunction (Halvorson et al., 2010). This pathology is characterized by fluid-filled cysts that replace normal renal parenchyma (Fig 7A). Aberrant HDAC6 activity has been implicated in cyst growth (Sun et al., 2019). Therefore, we reasoned that pharmacologically targeting HDAC6 may have therapeutic benefit in the reduction of cyst formation in cellular models of ADPKD (Fig 7B). Human pluripotent stem cells (hPSCs) with a targeted disruption of *PKD2* were differentiated into kidney organoids (Freedman et al., 2015). Cysts were identified as large, translucent structures that swayed in response to agitation (Fig 7C & D). *PKD2*^-/-^ kidney organoids and matched isogenic controls were treated with tubacin (0.2- 1µM) for 48 h and cyst size was evaluated. At low concentrations of tubacin (0.2 µM), renal cyst size was markedly reduced in *PKD2*^-/-^ organoids as compared to the control (Fig 7E & F). Similar results were obtained at a higher dose of 1µM (Fig 7G & H). Amalgamated data from five experiments are presented (Fig 7I). Drug toxicity as assessed by luminescence assay was evident at higher doses of tubacin (Fig 7J). Thus, pharmacologically targeting the AKAP220-binding partner HDAC6 reduces cystogenesis in a disease relevant model of ADPKD.

**Figure 7:**
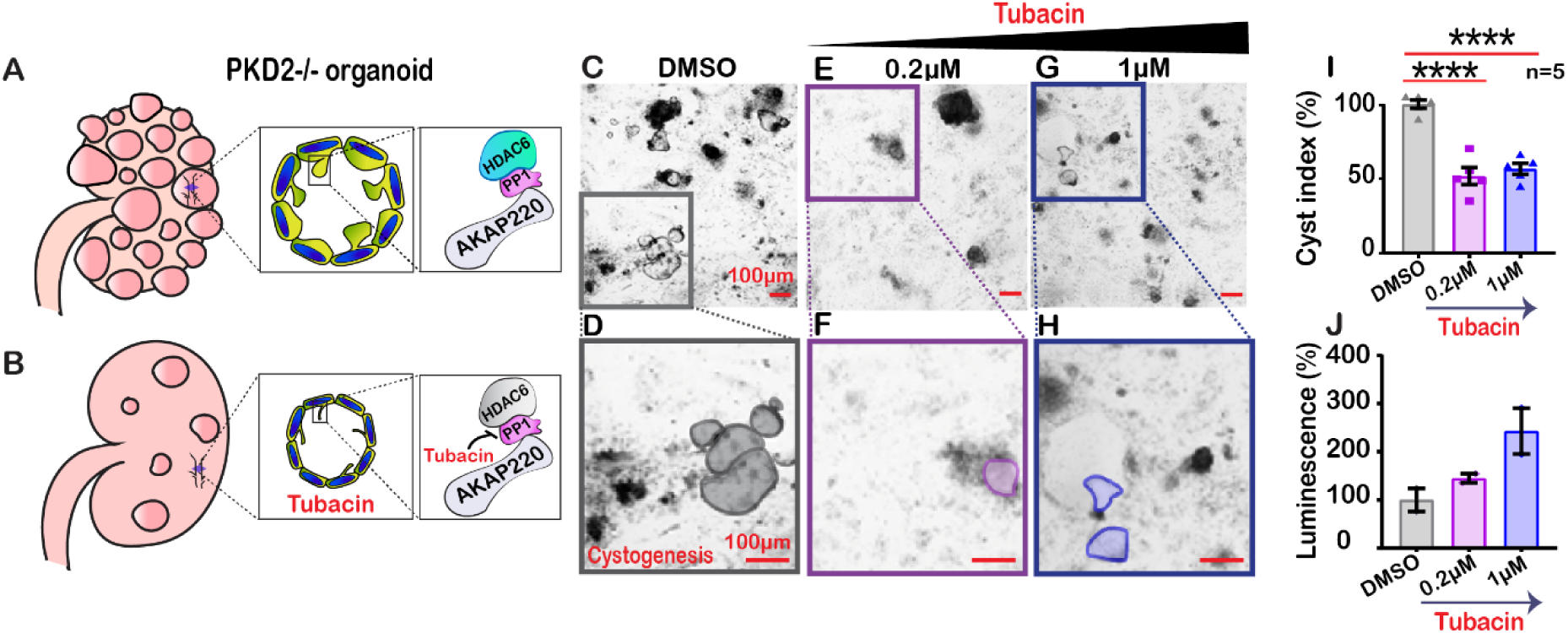
Inactivating HDAC6 reduces renal cystogenesis. Schematic of polycystic kidneys **A**) before and **B**) after tubacin treatment. **Insets**) Tubacin action on the AKAP220-signaling complex in precystic PKD2^-/-^ organoids derived from WA-09 cells. **C-J**) Confocal images of PKD2^-/-^ organoids treated with **C**) DMSO, **E**) 0.2 µM and **G**) 1µM tubacin. Enlarged regions from **C, E** and **G** showing cysts in **D**) DMSO, **F**) 0.2 µM and **H**) 1µM tubacin-treated conditions. Scale bars (100µm). **I**) Quantification (% cyst index) in DMSO (grey), 0.2 µM (purple) and 1µM (blue) tubacin-treated conditions. ****p<0.0001, N=5. **J**) Luminescence assay to detect toxicity of the drug is plotted for DMSO (grey), 0.2 µM (purple) and 1µM (blue) tubacin-treated conditions. Error bars are s.e.m. *P* values were calculated by unpaired two-tailed Student’s t-test.

## Discussion

The primary cilium is a highly organized mechanosensory transduction unit that responds to environmental cues (Goetz and Anderson, 2010; Wheway et al., 2018). We have discovered signaling events proceeding through AKAP220 that temper cilia development in kidney collecting ducts. While it may seem paradoxical that disruption of local signaling can positively impact organellar development, it is important to note that a few AKAPs mitigate signaling events in other cellular contexts (Bucko and Scott, 2021; Langeberg and Scott, 2015). At neuronal synapses, tonic phosphatase activity constrained by AKAP79/150 attenuates the phosphorylation status and activity of excitatory ionotropic glutamate receptors that contribute to learning and memory (Hoshi et al., 2005; Tunquist et al., 2008). In cardiomyocytes, AKAP18 anchored phosphodiesterase-3 degrades cAMP that sustains excitation-contraction coupling (Lygren et al., 2007). Likewise, AKAP220 binding partners repress aquaporin-2 shuttling to apical membranes of kidney collecting ducts to maintain renal water homeostasis (Whiting et al., 2016).

Our imaging analysis of tissue sections from AKAP220^-/-^ mice and cell lines indicate that loss of this anchoring protein enhances cilia development. Interestingly, AKAP79/150 has been implicated as a modulator of renal ciliary signaling (Choi et al., 2011). However, proximity proteomic approaches identify AKAP220 in cilia, and knockout of the AKAP79/150 ortholog in mIMCD3 cells and mice does not impact this aspect of cilia development ((May et al., 2020) & Supp Fig 1). In contrast, our *in vivo* and *in vitro* data strongly implicate AKAP220 signaling in the modulation of aquaporin-2 trafficking and ciliogenesis. This could occur through one of two mechanisms- either the cellular signals that attenuate aquaporin-2 trafficking and diminish cilia development are processed through the same macromolecular complex, or, distinct signaling islands of AKAP220-binding partners are assembled to control each process.

A unifying principle of our study is that AKAP220-binding partners affect the development of primary cilia by enacting cytoskeletal changes at the level of actin polymerization and tubulin acetylation. This implicates deacetylation as a fundamental signal termination process that checks the rate of cilia development. Histone deacetylase 6 (HDAC6) that targets the cytoskeletal elements alpha tubulin and cortactin has been identified as a driver of cilia disassembly (Ran et al., 2015). In concordance with this notion, data in figure 2Q show that rescue upon overexpression of HDAC6 in an AKAP220KO background restores control of ciliation. Several lines of evidence suggest a role for AKAP220 in this process. Data in figure 2 indicate that this deacetylase is more labile in the absence of the anchoring protein. While phosphorylation protects HDAC6 from ubiquitination, and dephosphorylation favors proteasomal degradation, it has been unclear how this enzyme is maintained in proximity of developing cilia (Ran et al., 2020). Our findings point towards a previously unrecognized adaptor function for protein phosphatase 1 (PP1). This reasoning is predicated on evidence that AKAP220 is a conventional PP1-targeting subunit that utilizes a KVxF motif to contact the phosphatase (Bollen et al., 2010).

While the mechanism of HDAC6 interaction with PP1 is less clear (Brush et al., 2004), it appears that the anchored phosphatase retains the capacity to function as an adaptor protein and recruit this additional binding partner. This is inferred by pulse-chase data showing that HDAC6 protein stability is compromised in AKAP220KO cells and supported by proximity-ligation data detecting HDAC6 in AKAP220-signaling islands (Fig 2I-O). Thus, AKAP220-PP1 subcomplexes create a platform for the targeting of HDAC6 to repress tubulin acetylation during ciliogenesis. This may represent a homeostatic mechanism that enables cells to enter mitosis.

Mutations in the ciliary phosphoproteins polycystin 1 and 2 are linked to autosomal dominant polycystic kidney disease (ADPKD) (Hughes et al., 1995; Mochizuki et al., 1996; Streets and Ong, 2020). Trafficking of polycystin-1 to the ciliary membrane is dependent on local PP1 activity (Parnell et al., 2012). Likewise, bi-directional control of polycystin-2 channel conductance is modulated by an anchored PKA-PP1 component (Streets et al., 2013). Thus, local dephosphorylation events are key to the termination of mechanotransduction signals that govern primary cilia action in ADPKD. Using the AKAP220-ΔPP1 knock-in mutant as a mechanistic probe, we have uncovered a new non-catalytic role for anchored-PP1 in the regulation of cytoskeletal events that underlie renal cilia biogenesis. The striking changes observed in actin remodeling and cilia morphology presented in figures 4 and 5 can solely be attributed to disruption of the KVxF motif in AKAP220 (Schillace and Scott, 1999). Concomitant effects on HDAC6 location not only enhance cortactin acetylation, but also enact changes in actin dynamics.

Such cytoskeletal reorganization at defined sites is a prelude to cilia extension. Our data in figures 6F-N indicate that these effects appear to be more pronounced in live cell experiments and within the pseudo physiological environment of the kidney-on-a-chip. Focal adhesion proteins attach the basal body to actin filaments (Drummond et al., 2018; Kim et al., 2015). This signals clearing of the local actin barrier to commence microtubule nucleation. In keeping with this notion, data in figure 5 show that the actin depolymerizing drug Cytochalasin D enhances cilia persistence and length, whereas the actin-stabilizing compound Jasplakinolide has the opposite effect (Casella et al., 1981; Holzinger, 2009). These pharmacological tools highlight the action of AKAP220- associated enzymes at the level of the actin barrier. This substructure is part of a “ciliary necklace” that acts as a physical checkpoint for proteins moving into the cilium (Long and Huang, 2020). This is supported by photobleaching studies showing that Arl13b moves more readily in AKAP220-ΔPP1 mutant cilia and these organelles are approximately 2-fold longer than wildtype (Fig 6H). Thus, it is plausible that AKAP220-associated PP1 participates in maintenance of an actin barrier close to the basal body of the cilium, the loss of which leads to unhindered protein movement into the developing organelle.

Understanding the signaling mechanisms that govern primary cilia biogenesis in collecting ducts has potential for the treatment of autosomal dominant polycystic kidney disease (ADPKD). Our studies provide the first demonstration of tubacin drug action in a human organoid model system for ADPKD. Our work points towards the AKAP220-PP1-HDAC6 signaling axis as a therapeutic target for this ciliopathy. Two lines of evidence converge on this notion. First, HDAC6 mediated deacetylation of tubulin controls cilia depolymerization. HDAC6 inhibitors including tubastatin A, ACY-1215 and tubacin are in early phase clinical trials to manage cancers and ciliopathies (Dong et al., 2018; Haggarty et al., 2003; Song et al., 2020). Our pharmacological studies now utilize tubacin to enhance ciliogenesis in wildtype mIMCD3 cells. Importantly, we show in figure 3Y that this selective-HDAC6 inhibitor has no effect in AKAP220-ΔPP1 cells. This argues that the AKAP220-PP1-HDAC6 axis is a molecular target for this drug. Consequently, tubacin could be exploited as a precision pharmaceutical to enhance ciliogenesis in disorders that arise from loss of primary cilia. Second, since HDAC6 activity is elevated in *PKD1* mutant renal epithelial cells (Li et al., 2016), we examined tubacin action in human cellular models of ADPKD. In figure 7, we show that tubacin reduced cystogenesis in *PKD2^-/-^* precystic organoids. Hence, targeting the AKAP220-PP1-HDAC6 subcomplex may not only enhance cilia development, but also interfere with downstream signaling events that contribute to cystogenesis. The AKAP-targeting concept has recently been used to restrain kinase inhibitor drugs at defined subcellular locations (Bucko et al., 2019, 2020). We will expand this approach towards developing a precision pharmacology strategy to selectively deliver HDAC6 inhibitors to primary cilia for the treatment of chronic disorders such as polycystic kidney disease.

## Materials and methods

### Tissue section immunofluorescent staining

Kidneys were fixed in 10% (vol/vol) buffered formalin (4 °C), embedded in paraffin and 4-μm-thick sections collected. Sections were deparaffinized using Citrasolv (Fisher) and antigen retrieved in buffer A using a Retriever 2100 pressure cooker (Electron Microscopy Sciences). Tissue sections were blocked in 10% (vol/vol) donkey serum in PBS solution before overnight incubation with the respective primary antibodies.

### mIMCD3 spheroids

mIMCD3 cells were seeded in Matrigel to generate spheroids as previously described (Giles et al., 2014). The spheroids were stained with Acetyl tubulin for marking primary cilia and those with a visible open lumen were imaged and used in the quantification.

### Cell culture

mIMCD3 cells were maintained in DMEM: F12 1:1 media supplemented with 10% FBS and penicillin/streptomycin. Cells were transfected with plasmids using Mirus TransIT-LT1 transfection reagent and incubated for 48-72 hrs before lysis or fixation. All cell lines were maintained in a 5% CO2 incubator at 37 °C.

### Virus generation

Constitutively active lentiviral plasmids expressing Arl13b-GFP were transfected into HEK cells along with viral packaging and envelope plasmids. The viral particles generated are filtered and added to mIMCD3 cells. After 24 hours of incubation, the cells were selected in zeocin (400ug/ml) for the next 2 weeks. After selection, the cells were trypsinized and plated at a low density of about 1 cell/well in 96 well plate, expanded and tested for expression of Arl13b-GFP by western blotting and immunofluorescence.

### Immunoblotting and blot analysis

Cells were grown to the desired confluence and washed once with PBS at room temperature. Cold lysis buffer (20 mM HEPES, (pH 7.4),150 mM NaCl,1 mM EDTA,1% triton X-100 in water) was added along with protease and phosphatase inhibitors and the plate was rocked gently at 4 degrees for 10 mins. The cell lysate was then scarped into a pre-chilled tube and cleared at 12,000*g for 10 min at 4 degrees. A BCA assay (Pierce) was used to determine protein concentrations, and 30µg of protein was loaded onto a Bolt 4–12% bis-Tris gel (Life Technologies). The cleared lysate was boiled in 2X SDS loading buffer for 10 min before loading. Proteins were transferred to nitrocellulose membrane and blocked in either 5% milk. The blot was incubated in primary antibody at 1:1000 or as specified by the manufacturer overnight at 4 °C. Immunoblots were washed (3 times, 10 mins each) in TBST before incubation in a 1:10,000 secondary antibody for 1 hr at room temperature. Immunoblots were washed again in TBST (3 times, 10 mins each) before imaging on an iBright FL1000 (Thermo Fisher Scientific) with SuperSignal Dura ECL reagent (Thermo Fisher Scientific). Densitometry for blot quantification was done using thermo fisher’s software.

### Immunofluorescence and microscopy

#### Sample preparation

Cells were seeded on acid-washed coverslips in 12-well plates. After they achieve the desired confluence, the wells were rinsed thrice with PBS and fixed with 4% paraformaldehyde in PBS or 10% ice cold methanol (based on the antibody specification) for 12 mins. After fixation, the cells were permeabilized in PBS+0.1%Triton x-100+1%BSA for 12 mins, blocked with 2%BSA for 30 mins and treated with the respective primary antibodies overnight. After thorough washing in PBS the next day (3 times, 5mins each), secondary antibodies conjugated to Alexa fluor dyes were added for 2 hours. After 2 washes in PBS, DAPI along with or without the actin probe (based on the experiment) was added for 10 mins. The coverslips were washed 2 more times in PBS and mounted on slides using ProLong Diamond Antifade Mountant (Life Technologies).

#### Imaging and analysis

Cells were imaged on a Keyence BZ-X710 microscope (Keyence, Itasca, IL) using the relevant filter cubes for DAPI (blue filter), Actin (red filter), Cortactin (green filter). All images were acquired with the same magnification (100X, oil immersion), exposure time, and illumination intensity. Images were quantified and processed using ImageJ software.

### FRAP (Fluorescence recovery after photobleaching)

#### Sample preparation

Cells were reverse transfected with lifeact-GFP (Addgene plasmid no.58470) and seeded into 60mm glass-bottom plates for 24 hours. The transfected plates are rinsed in PBS and incubated in Fluorobrite medium containing NucBlue Hoescht 33342 stain (R37605, Invitrogen, 1 drop/ml). For Arl13b FRAP, mIMCD3 lines were transduced with L13- Arl13bGFP lentiviral construct and selected as outlined above. These cells were seeded in 60mm glass-bottom plates. They were treated with 0.1%FBS for 24 hours and then the experiment was performed.

#### Imaging technique and analysis

Cells were imaged on a GE Deltavision OMX SR microscope (GE Life Healthcare Sciences). After loading (as above), cells were placed in a humidified chamber with 5% CO2 at 37 °C and imaged using a 60X oil immersion objective (Olympus, Shinjuku, Tokyo, Japan). The actin signal was photobleached in the green channel (488nm) using a 15% laser power for 0.05 seconds. A total of 5 events were captured before photobleaching. Images after photobleaching were captured every 100 milliseconds for 10 seconds. For Arl13b FRAP, Lentiviral Arl13b GFP mIMCD3 cells. The Laser pulse at 488 was at 30% T, in a spot at the tip of the cilium to bleach it for 0.1 seconds. 5 events before the bleach and a total of 350 time points were taken at 10msec increments. All images were acquired with the same settings. Images were quantified and processed using ImageJ software.

### Antibodies

The following antibodies were used in this study for immunoblotting: AKAP220 (custom rabbit polyclonal,1:1000), GAPDH (Novus, mouse monoclonal, 1:5000), Acetylated tubulin (Thermo Fisher # 32-2700, 1:1000), HDAC6 (Proteintech # 12834-1-AP, 1:1000). Antibodies used for immunofluorescence: Acetylated tubulin (Thermo Fisher # 32-2700, 1:10000), HDAC6 (Proteintech, 12834-1-AP, 1:400), Arl13b (Proteintech # 17711-1-AP, 1:5000), Actin (Molecular probe, R37112, 1drop/ml).

### Plasmids

Lentiviral Arl13b-GFP (# 40879) and lifeact-GFP (#58470) were purchased from addgene.

### Statistics for all experiments

Statistical analysis was performed using an unpaired two-tailed Student’s t-test or one-way ANOVA (based on the number of samples) in GraphPad Prism software. All values are reported as mean ± standard error of the mean (s.e.m) with p-values less than 0.05 considered statistically significant. For each experiment, number of independent experiments (N) and number of individual points from all experiments (n) are presented.

### Line-plot analysis in ImageJ

For the actin and cortactin distribution experiments, lines were drawn from the outer edge of the nucleus to beyond the lamellipodia of the cell and line plots were generated using ImageJ to give the pixel intensity of each of the proteins along the line.

### HDAC6 activity assay

The experiment was performed using BioVision’s HDAC6 activity assay kit. mIMCD3 cells were seeded in 10cm dishes and allowed to grow for the desired confluence. They were then lysed using the lysis buffer provided in the kit. The protein concentration was measured using BCA assay kit (Pierce) and 20µg of protein was loaded per well. 5µM tubacin was used in the experiment and the fluorescence intensity was measured at 380/490 nm in the plate reader.

### Drug treatment experiments

mIMCD3 cells were seeded on cover slips in 12-well plates and allowed to grow to the desired confluence. They were treated with 2uM tubacin for 4 hours, 200nM cytochalasin D for 4 hours or 500nM Jasplakinolide for 1.5 hours and fixed immediately after treatment (as previously described). The cells were stained for acetyl tubulin and Arl13b to mark primary cilia and DAPI to stain DNA.

### Organoid differentiation

Organoids were differentiated in 384 well plates from human pluripotent stem cells (WTC11 iPS cells, Conklin lab, Gladstone Institute) that had been modified to disrupt *PKD2* (Cruz et al., 2017).

## Acknowledgments

We thank Scott Lab members for their valuable feedback on the manuscript. This work was supported by NIH: 5R01DK105542 and 1R01DK119192-01 (JDS), NIH Awards T32 GM007270 (Gopalan), UG3TR002158 (Himmelfarb) and R01DK117914 (Freedman), F32DK121415 (Omar) and the Lara Nowak Macklin Research Fund.

## Author contributions

J.G. and J.D.S. conceived and designed the study. J.G. designed experiments. J.G., M.H.O., A.R. and J.F. performed experiments. J.G, K.F and N.C made critical reagents for the study. J.G., M.H.O., A.R. and J.F. analyzed the data. J.D.S. and B.S.F. provided critical advice on experimental design, data analyses, and data interpretations. J.G. and J.D.S. prepared figures and wrote the manuscript.

## Data availability

All raw images and data can be provided upon request.

---

**Supplemental figure 1:**
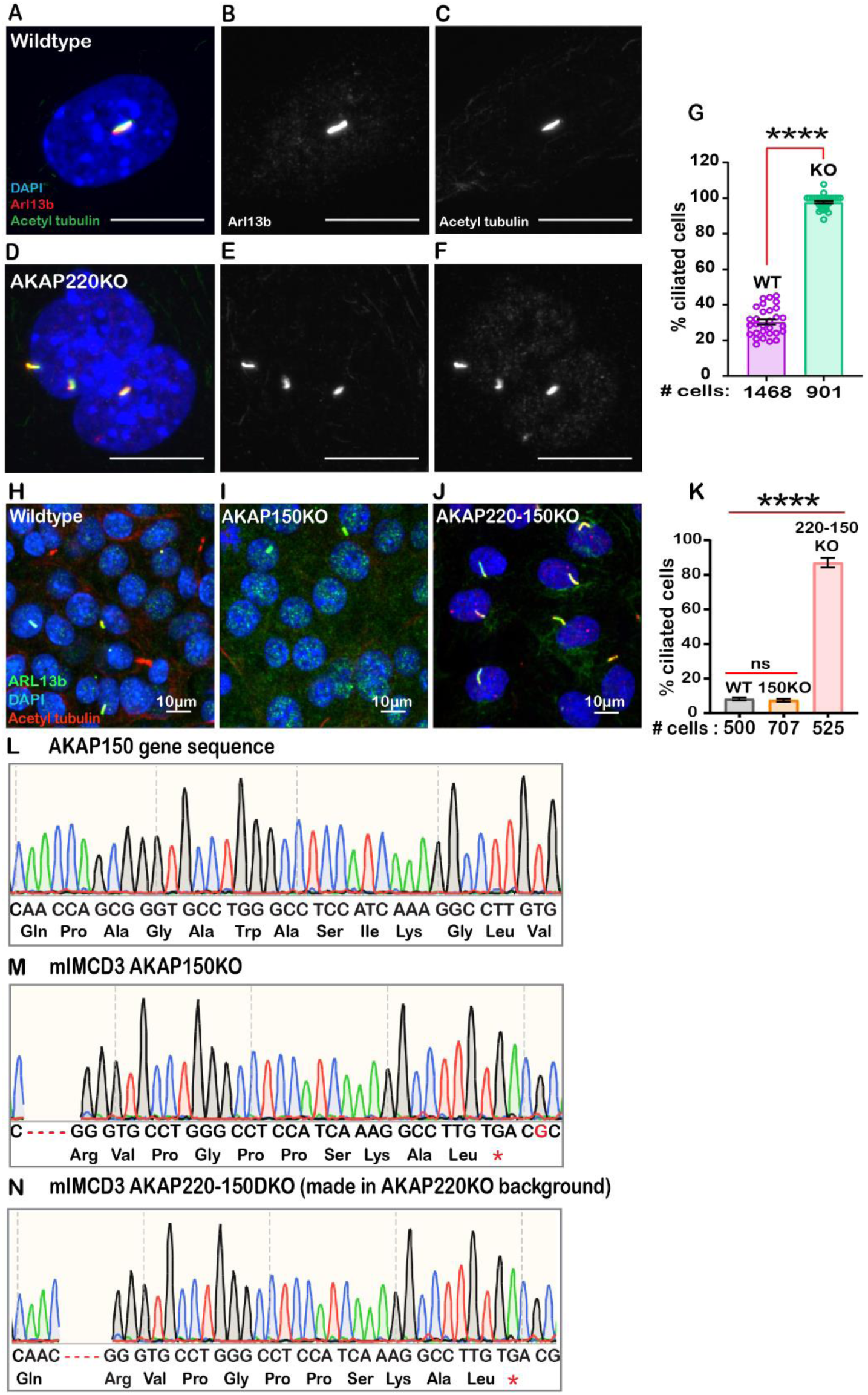
Quantification of cilia number in gene-edited mIMCD3 cells. Immunofluorescent staining of Arl13b (red), acetyl tubulin (green) and DAPI (blue) in serum starved (0.5% FBS, 24hrs) **A**) wildtype and **D**) AKAP220KO mIMCD3 cells. Grey scale images of Arl13b (**B & E**) and acetyl tubulin (**C & F**) show multiple cilia in the AKAP220KO cell. **G**) Quantification (% ciliated cells) in wildtype (purple column), AKAP220KO (green column). ****p<0.0001. Number of cells analyzed are indicated below each bar. **H-K) Deletion of AKAP150 has no effect on primary cilia development**. Crispr-Cas 9 gene editing was used to delete the murine anchoring protein AKAP150 in mIMCD3 cells. Double knockout cells were also produced lacking AKAP220 and AKAP150. Immunofluorescent staining with ciliary markers Arl13b (green) and acetyl tubulin (red) in **H**) wildtype, **I**) AKAP150KO and **J**) AKAP220-150KO mIMCD3 cells. DAPI serves as a nuclear marker. **K**) Quantification (% ciliated cells) in wildtype (grey column), AKAP150KO (orange column) and AKAP220-150KO (coral column). Number of cells analyzed are indicated below each bar. ****p<0.0001, ns=non-significant, N=3. All error bars are s.e.m. *P* values were calculated by unpaired two-tailed Student’s t-test. Scale bars (10µm). CRISPR-Cas9 gene editing was used to disrupt the AKAP150 gene to generate AKAP150 and AKAP220-150 double knockout mIMCD3 cells. Sequencing analysis data shows **L**) intact AKAP150 in wildtype AKAP150, and deletions of AKAP150 in **M**) AKAP150KO cells and **N**) AKAP220-150 double knockout cells made in AKAP220KO background.

**Supplemental figure 2:**
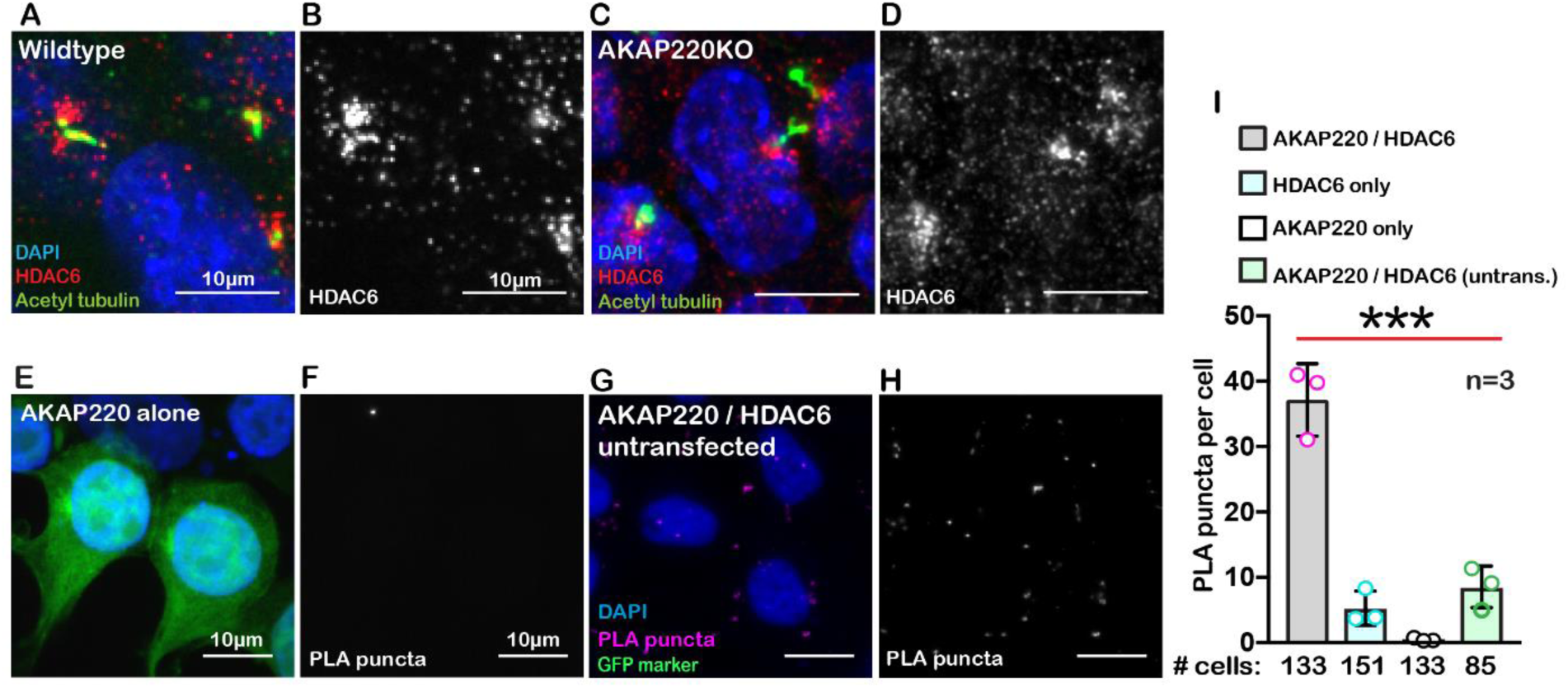
Further characterization of AKAP220-HDAC6 interface. HDAC6 accumulates at the base of primary cilia. Immunofluorescent detection of acetyl tubulin (green), HDAC6 (red) and DAPI (blue) in **A**) wildtype and **C**) AKAP220KO mIMCD3 cells. Grey scale images of HDAC6 in **B**) wildtype and **D**) AKAP220KO show localization of the deacetylase at the base of primary cilia. **Additional Proximity ligation (PLA) controls**. Proximity ligation (PLA) detection of V5-AKAP220/HDAC6 subcomplexes (pink), DAPI (blue) in cells expressing GFP (green) as a transfection marker in mIMCD3 cells **E**) transfected with V5-AKAP220 only and **G**) AKAP220/HDAC6 untransfected. **F**) and **H**) Grey scale image highlights V5- AKAP220/HDAC6 PLA puncta. **I**) Amalgamated data (PLA puncta/cell) from three independent experiments is presented.

**Supplemental figure 3:**
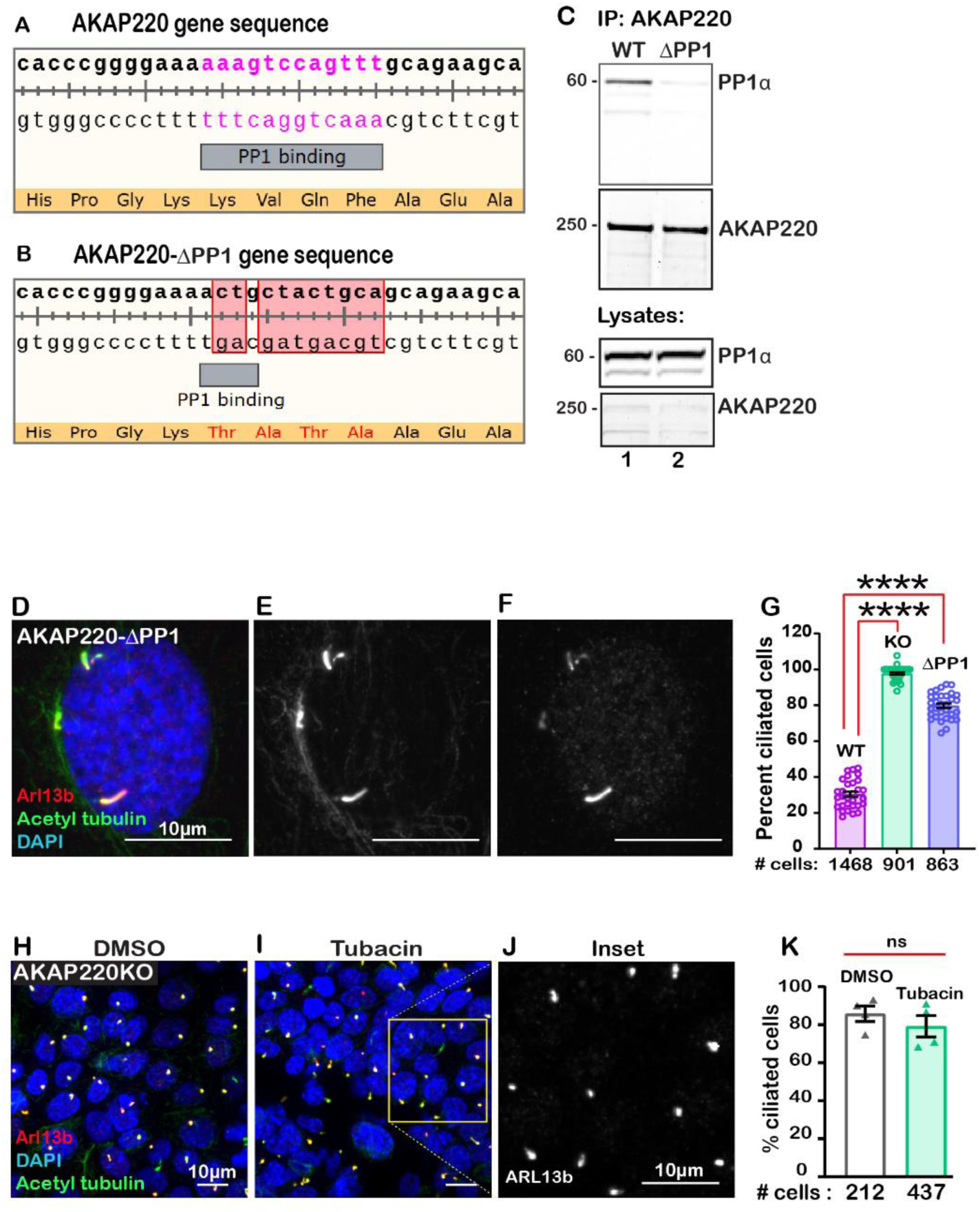
Further characterization of AKAP220-ΔPP1 and tubacin action. CRISPR-Cas9 gene editing was used to substitute the PP1-targeting KVQF motif in AKAP220 gene to generate AKAP220-ΔPP1 mIMCD3 cells. Sequencing analysis data shows **A**) wildtype PP1-binding region in AKAP220 and **B**) modified TATA region in AKAP220-ΔPP1. **C**) AKAP220- ΔPP1 does not recruit PP1. Immunoblot detection shows PP1 (top panel) in WT AKAP220 immune complexes (lane 1), but not in AKAP220-ΔPP1 immune complexes (lane 2). **D-F) Immunofluorescent** staining of Arl13b (red), acetyl tubulin (green) and DAPI (blue) in serum starved (0.5% FBS, 24hrs). **D**) AKAP220-ΔPP1 mIMCD3 cells. Grey scale images of **E**) Arl13b and **F**) acetyl tubulin show multiple cilia in the AKAP220-ΔPP1 cell. **G**) Quantification (% ciliated cells) in wildtype (purple column), AKAP220KO (green column) and AKAP220-ΔPP1 (blue column). ****p<0.0001. Number of cells analyzed are indicated below each bar. Immunofluorescent staining with ciliary markers acetyl tubulin (green) and Arl13b (red) in **H**) DMSO and **I**) tubacin-treated AKAP220KO cells. DAPI (blue) serves as a nuclear marker. **J**) Expanded grey scale image of Arl13b in tubacin-treated AKAP220KO cells. **K**) Quantification (% ciliated cells) in DMSO (black bar) and tubacin-treated (green bar) AKAP220KO cells. The number of cells analyzed from three independent experiments are indicated below the bar for each condition. ns=non-significant. Error bars are s.e.m. *P* values were calculated by unpaired two-tailed Student’s t-test. Scale bars (10µm).

**Supplemental figure 4:**
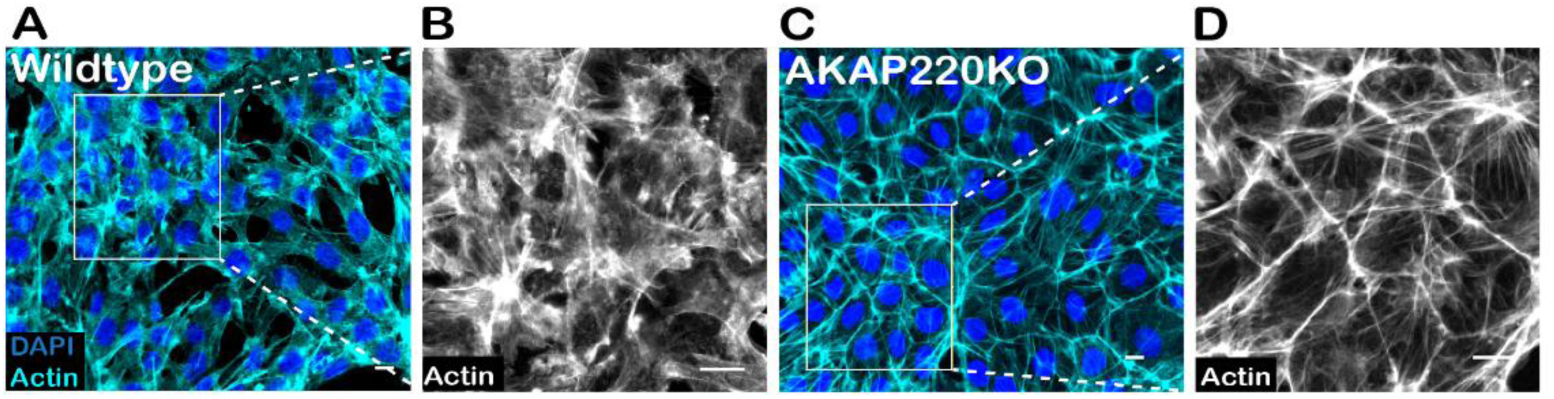
Distribution of actin in confluent mIMCD3 cell culture. Immunofluorescent detection of actin (cyan) and DAPI (blue) in A) wildtype and C) AKAP220KO mIMCD3 cells. Grey scale images of actin in B) wildtype and D) AKAP220KO cells show the distribution of actin in confluent cells. Scale bars (10µm).

**Supplemental figure 5:**
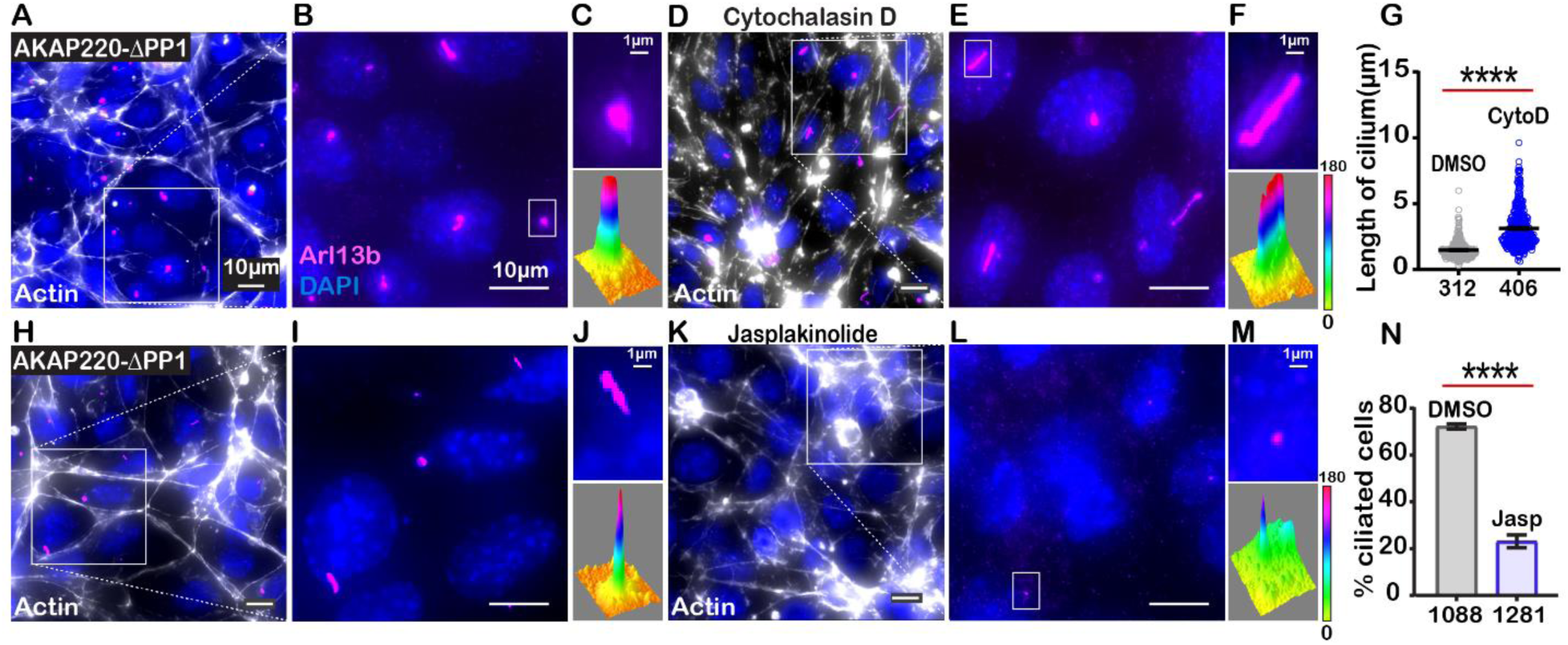
Characterizing the effect of actin-modulating drugs on AKAP220- ΔPP1 cilia. A-G) Immunofluorescent detection of actin (white), Arl13b (pink) and DAPI (blue) in A) DMSO and D) 200nM Cytochalasin D-treated AKAP220-ΔPP1 mIMCD3 cells. Inset) Expanded field of cells treated with B) DMSO or E) Cytochalasin D. Boxed regions in C and F focus on a single cilium (top) and three-dimensional surface plot (bottom). The width of the cylindrical region in the 3D surface plot represents cilium length. G) Quantification of cilia length in DMSO (grey) and Cytochalasin D (pink) treated cells. ****p<0.0001, N=3. H-N) Immunofluorescent staining of actin (white) and DAPI (blue) of H (DMSO) and K (Jasplakinolide) treated AKAP220-ΔPP1 cells. Inset) Expanded field of cells treated with I) DMSO or L) Jasplakinolide. Boxed regions in J and M focus on a single cilium (top) and three-dimensional surface plot (bottom). The width of the cylindrical region in the 3D surface plot represents cilium length. N) Quantification (% ciliated cells) in DMSO (grey) and Jasplakinolide (purple). ****p<0.0001, N=3. All error bars are s.e.m. *P* values were calculated by unpaired two-tailed Student’s t-test. Scale bars (10µm). Number of cells analyzed indicated below each column.

**Supplemental figure 6:**
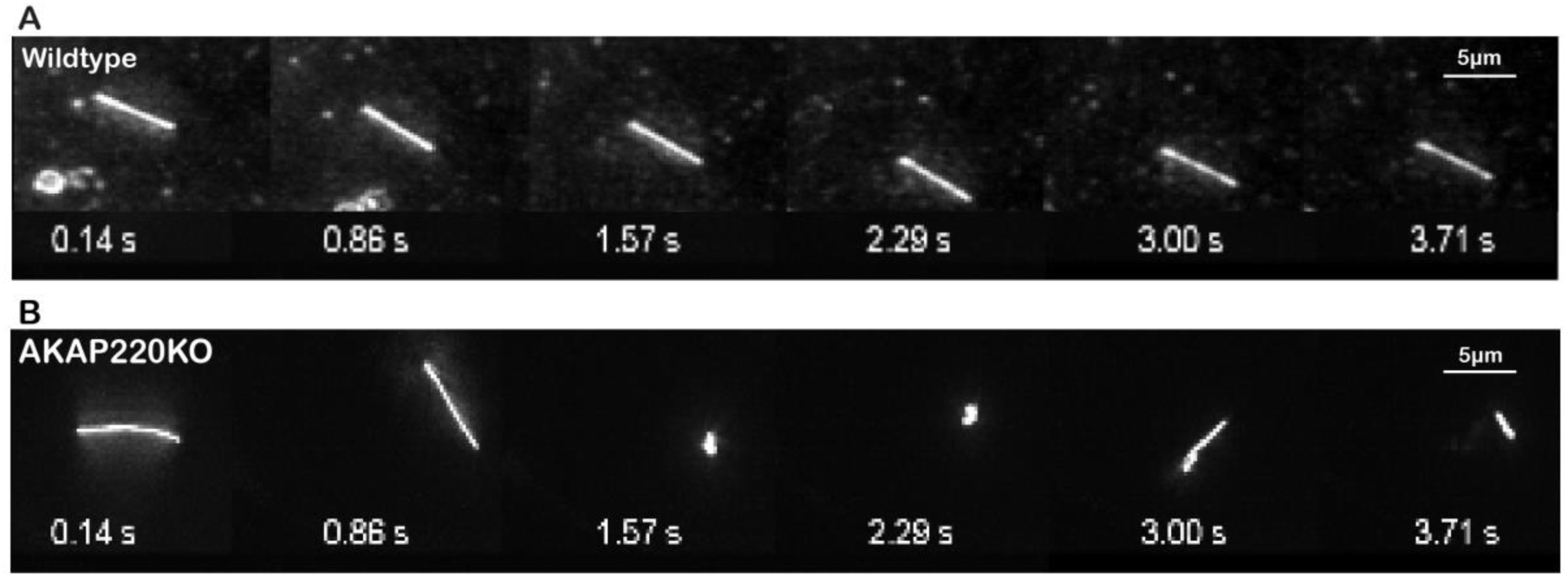
Super resolution videos depicting flexibility of AKAP220KO cilia. A) Wildtype and B) AKAP220KO primary cilia from mIMCD3 cells transduced with Arl13b-GFP were monitored across a time course of 4 seconds. The AKAP220 cilium is longer and more flexible compared to the wildtype.

## References

1. Badano, J.L., Mitsuma, N., Beales, P.L., and Katsanis, N. (2006). The Ciliopathies: An Emerging Class of Human Genetic Disorders. Annu. Rev. Genomics Hum. Genet. 7, 125–148.

2. Bankir, L., Bouby, N., and Ritz, E. (2013). Vasopressin: a novel target for the prevention and retardation of kidney disease? Nat. Rev. Nephrol. 9, 223–239.

3. Bastos, A.P., and Onuchic, L.F. (2011). Molecular and cellular pathogenesis of autosomal dominant polycystic kidney disease. Braz. J. Med. Biol. Res. Rev. Bras. Pesqui. Medicas E Biol. 44, 606–617.

4. Blattner, S.M., Hodgin, J.B., Nishio, M., Wylie, S.A., Saha, J., Soofi, A.A., Vining, C., Randolph, A., Herbach, N., Wanke, R., et al. (2013). Divergent functions of the Rho GTPases Rac1 and Cdc42 in podocyte injury. Kidney Int. 84, 920–930.

5. Bollen, M., Peti, W., Ragusa, M.J., and Beullens, M. (2010). The extended PP1 toolkit: designed to create specificity. Trends Biochem. Sci. 35, 450–458.

6. Breslow, D.K., Hoogendoorn, S., Kopp, A.R., Morgens, D.W., Vu, B.K., Kennedy, M.C., Han, K., Li, A., Hess, G.T., Bassik, M.C., et al. (2018). A CRISPR-based screen for Hedgehog signaling provides insights into ciliary function and ciliopathies. Nat. Genet. 50, 460–471.

7. Brush, M.H., Guardiola, A., Connor, J.H., Yao, T.-P., and Shenolikar, S. (2004). Deactylase inhibitors disrupt cellular complexes containing protein phosphatases and deacetylases. J. Biol. Chem. 279, 7685–7691.

8. Bucko, P.J., and Scott, J.D. (2021). Drugs that Regulate Local Cell Signaling: AKAP Targeting as a Therapeutic Option. Annu. Rev. Pharmacol. Toxicol. 61, null.

9. Bucko, P.J., Lombard, C.K., Rathbun, L., Garcia, I., Bhat, A., Wordeman, L., Smith, F.D., Maly, D.J., Hehnly, H., and Scott, J.D. (2019). Subcellular drug targeting illuminates local kinase action. ELife 8, e52220.

10. Bucko, P.J., Garcia, I., Manocha, R., Bhat, A., Wordeman, L., and Scott, J.D. (2020). Gravin-associated kinase signaling networks coordinate γ-tubulin organization at mitotic spindle poles. J. Biol. Chem. jbc.RA120.014791.

11. Casella, J.F., Flanagan, M.D., and Lin, S. (1981). Cytochalasin D inhibits actin polymerization and induces depolymerization of actin filaments formed during platelet shape change. Nature 293, 302–305.

12. Choi, Y.-H., Suzuki, A., Hajarnis, S., Ma, Z., Chapin, H.C., Caplan, M.J., Pontoglio, M., Somlo, S., and Igarashi, P. (2011). Polycystin-2 and phosphodiesterase 4C are components of a ciliary A-kinase anchoring protein complex that is disrupted in cystic kidney diseases. Proc. Natl. Acad. Sci. 108, 10679–10684.

13. Dong, J., Zheng, N., Wang, X., Tang, C., Yan, P., Zhou, H.-B., and Huang, J. (2018). A novel HDAC6 inhibitor exerts an anti-cancer effect by triggering cell cycle arrest and apoptosis in gastric cancer. Eur. J. Pharmacol. 828, 67–79.

14. Drummond, M.L., Li, M., Tarapore, E., Nguyen, T.T.L., Barouni, B.J., Cruz, S., Tan, K.C., Oro, A.E., and Atwood, S.X. (2018). Actin polymerization controls cilia-mediated signaling. J. Cell Biol. 217, 3255–3266.

15. Farina, F., Gaillard, J., Guérin, C., Couté, Y., Sillibourne, J., Blanchoin, L., and Théry, M. (2016). The centrosome is an actin-organizing centre. Nat. Cell Biol. 18, 65–75.

16. Fliegauf, M., Benzing, T., and Omran, H. (2007). When cilia go bad: cilia defects and ciliopathies. Nat. Rev. Mol. Cell Biol. 8, 880–893.

17. Freedman, B.S., Lam, A.Q., Sundsbak, J.L., Iatrino, R., Su, X., Koon, S.J., Wu, M., Daheron, L., Harris, P.C., Zhou, J., et al. (2013). Reduced ciliary polycystin-2 in induced pluripotent stem cells from polycystic kidney disease patients with PKD1 mutations. J. Am. Soc. Nephrol. JASN 24, 1571–1586.

18. Freedman, B.S., Brooks, C.R., Lam, A.Q., Fu, H., Morizane, R., Agrawal, V., Saad, A.F., Li, M.K., Hughes, M.R., Werff, R.V., et al. (2015). Modelling kidney disease with CRISPR-mutant kidney organoids derived from human pluripotent epiblast spheroids. Nat. Commun. 6, 8715.

19. Giles, R.H., Ajzenberg, H., and Jackson, P.K. (2014). 3D spheroid model of mIMCD3 cells for studying ciliopathies and renal epithelial disorders. Nat. Protoc. 9, 2725–2731.

20. Goetz, S.C., and Anderson, K.V. (2010). The primary cilium: a signalling centre during vertebrate development. Nat. Rev. Genet. 11, 331–344.

21. Haggarty, S.J., Koeller, K.M., Wong, J.C., Grozinger, C.M., and Schreiber, S.L. (2003). Domain-selective small-molecule inhibitor of histone deacetylase 6 (HDAC6)-mediated tubulin deacetylation. Proc. Natl. Acad. Sci. U. S. A. 100, 4389–4394.

22. Halvorson, C.R., Bremmer, M.S., and Jacobs, S.C. (2010). Polycystic kidney disease: inheritance, pathophysiology, prognosis, and treatment. Int. J. Nephrol. Renov. Dis. 3, 69– 83.

23. Harris, P.C., and Torres, V.E. (2014). Genetic mechanisms and signaling pathways in autosomal dominant polycystic kidney disease. J. Clin. Invest. 124, 2315–2324.

24. Hemmelgarn, B.R., Zhang, J., Manns, B.J., Tonelli, M., Larsen, E., Ghali, W.A., Southern, D.A., McLaughlin, K., Mortis, G., and Culleton, B.F. (2006). Progression of kidney dysfunction in the community-dwelling elderly. Kidney Int. 69, 2155–2161.

25. Holzinger, A. (2009). Jasplakinolide: an actin-specific reagent that promotes actin polymerization. Methods Mol. Biol. Clifton NJ 586, 71–87.

26. Hoshi, N., Langeberg, L.K., and Scott, J.D. (2005). Distinct enzyme combinations in AKAP signalling complexes permit functional diversity. Nat. Cell Biol. 7, 1066–1073.

27. Hubbert, C., Guardiola, A., Shao, R., Kawaguchi, Y., Ito, A., Nixon, A., Yoshida, M., Wang, X.-F., and Yao, T.-P. (2002). HDAC6 is a microtubule-associated deacetylase. Nature 417, 455–458.

28. Hughes, J., Ward, C.J., Peral, B., Aspinwall, R., Clark, K., San Millán, J.L., Gamble, V., and Harris, P.C. (1995). The polycystic kidney disease 1 (PKD1) gene encodes a novel protein with multiple cell recognition domains. Nat. Genet. 10, 151–160.

29. Janke, C., and Magiera, M.M. (2020). The tubulin code and its role in controlling microtubule properties and functions. Nat. Rev. Mol. Cell Biol. 21, 307–326.

30. . Jo, I., Ward, D.T., Baum, M.A., Scott, J.D., Coghlan, V.M., Hammond, T.G., and Harris, H.W. (2001). AQP2 is a substrate for endogenous PP2B activity within an inner medullary AKAP-signaling complex. Am. J. Physiol.-Ren. Physiol. 281, F958–F965.

31. Kim, J., Jo, H., Hong, H., Kim, M.H., Kim, J.M., Lee, J.-K., Heo, W.D., and Kim, J. (2015). Actin remodelling factors control ciliogenesis by regulating YAP/TAZ activity and vesicle trafficking. Nat. Commun. 6, 6781.

32. Klymkowsky, M.W. (1999). Weaving a tangled web: the interconnected cytoskeleton. Nat. Cell Biol. 1, E121–E123.

33. Langeberg, L.K., and Scott, J.D. (2015). Signalling scaffolds and local organization of cellular behaviour. Nat. Rev. Mol. Cell Biol. 16, 232–244.

34. Li, X., Qi, N., Li, L., Wu, M., and Mei, C. (2016). Cytosolic HDAC6 is accumulated in cystic kidneys. Kidney Int. 90, 705.

35. Logue, J.S., and Scott, J.D. (2010). Organizing signal transduction through A-kinase Anchoring Proteins (AKAPs). FEBS J. 277, 4370–4375.

36. Logue, J.S., Whiting, J.L., Tunquist, B., Sacks, D.B., Langeberg, L.K., Wordeman, L., and Scott, J.D. (2011). AKAP220 Protein Organizes Signaling Elements That Impact Cell Migration. J. Biol. Chem. 286, 39269–39281.

37. Long, H., and Huang, K. (2020). Transport of Ciliary Membrane Proteins. Front. Cell Dev. Biol. 7.

38. Luo, C., Wu, M., Su, X., Yu, F., Brautigan, D.L., Chen, J., and Zhou, A.J. (2019). Protein phosphatase 1α interacts with a novel ciliary targeting sequence of polycystin-1 and regulates polycystin-1 trafficking. FASEB J. 33, 9945–9958.

39. Lygren, B., Carlson, C.R., Santamaria, K., Lissandron, V., McSorley, T., Litzenberg, J., Lorenz, D., Wiesner, B., Rosenthal, W., Zaccolo, M., et al. (2007). AKAP complex regulates Ca2+ re-uptake into heart sarcoplasmic reticulum. EMBO Rep. 8, 1061–1067.

40. Ma, M., Tian, X., Igarashi, P., Pazour, G.J., and Somlo, S. (2013). Loss of cilia suppresses cyst growth in genetic models of autosomal dominant polycystic kidney disease. Nat. Genet. 45, 1004–1012.

41. Ma, M., Gallagher, A.-R., and Somlo, S. (2017). Ciliary Mechanisms of Cyst Formation in Polycystic Kidney Disease. Cold Spring Harb. Perspect. Biol. 9.

42. . May, E.A., Kalocsay, M., D’Auriac, I.G., Gygi, S.P., Nachury, M.V., and Mick, D.U. (2020). Time-resolved proteomic profiling of the ciliary Hedgehog response reveals that GPR161 and PKA undergo regulated co-exit from cilia. BioRxiv 2020.07.29.225797.

43. Mochizuki, T., Wu, G., Hayashi, T., Xenophontos, S.L., Veldhuisen, B., Saris, J.J., Reynolds, D.M., Cai, Y., Gabow, P.A., Pierides, A., et al. (1996). PKD2, a gene for polycystic kidney disease that encodes an integral membrane protein. Science 272, 1339–1342.

44. . Mukhopadhyay, S., Wen, X., Ratti, N., Loktev, A., Rangell, L., Scales, S.J., and Jackson, P.K. (2013). The ciliary G-protein-coupled receptor Gpr161 negatively regulates the Sonic hedgehog pathway via cAMP signaling. Cell 152, 210–223.

45. Noda, Y., Sohara, E., Ohta, E., and Sasaki, S. (2010). Aquaporins in kidney pathophysiology. Nat. Rev. Nephrol. 6, 168–178.

46. Omar, M.H., and Scott, J.D. (2020). AKAP Signaling Islands: Venues for Precision Pharmacology. Trends Pharmacol. Sci. 41, 933–946.

47. Parnell, S.C., Puri, S., Wallace, D.P., and Calvet, J.P. (2012). Protein Phosphatase-1α Interacts with and Dephosphorylates Polycystin-1. PLoS ONE 7.

48. Portran, D., Schaedel, L., Xu, Z., Théry, M., and Nachury, M.V. (2017). Tubulin acetylation protects long-lived microtubules against mechanical ageing. Nat. Cell Biol. 19, 391–398.

49. Ran, J., Yang, Y., Li, D., Liu, M., and Zhou, J. (2015). Deacetylation of α-tubulin and cortactin is required for HDAC6 to trigger ciliary disassembly. Sci. Rep. 5, 12917.

50. Ran, J., Liu, M., Feng, J., Li, H., Ma, H., Song, T., Cao, Y., Zhou, P., Wu, Y., Yang, Y., et al. (2020). ASK1-Mediated Phosphorylation Blocks HDAC6 Ubiquitination and Degradation to Drive the Disassembly of Photoreceptor Connecting Cilia. Dev. Cell 53, 287–299.e5.

51. Saborio, P., Krieg Jr., R.J., Kuemmerle, N.B., Norkus, E.P., Schwartz, C.C., and Chan, J.C.M. (2000). α-Tocopherol modulates lipoprotein cytotoxicity in obstructive nephropathy. Pediatr. Nephrol. 14, 740–746.

52. Satir, P., Pedersen, L.B., and Christensen, S.T. (2010). The primary cilium at a glance. J. Cell Sci. 123, 499–503.

53. Schillace, R.V., and Scott, J.D. (1999). Association of the type 1 protein phosphatase PP1 with the A-kinase anchoring protein AKAP220. Curr. Biol. CB 9, 321–324.

54. Smith, F.D., Esseltine, J.L., Nygren, P.J., Veesler, D., Byrne, D.P., Vonderach, M., Strashnov, I., Eyers, C.E., Eyers, P.A., Langeberg, L.K., et al. (2017). Local protein kinase A action proceeds through intact holoenzymes. Science 356, 1288–1293.

55. Smith, F.D., Omar, M.H., Nygren, P.J., Soughayer, J., Hoshi, N., Lau, H.-T., Snyder, C.G., Branon, T.C., Ghosh, D., Langeberg, L.K., et al. (2018). Single nucleotide polymorphisms alter kinase anchoring and the subcellular targeting of A-kinase anchoring proteins. Proc. Natl. Acad. Sci. 115, E11465–E11474.

56. Somatilaka, B.N., Hwang, S.-H., Palicharla, V.R., White, K.A., Badgandi, H., Shelton, J.M., and Mukhopadhyay, S. (2020). Ankmy2 Prevents Smoothened-Independent Hyperactivation of the Hedgehog Pathway via Cilia-Regulated Adenylyl Cyclase Signaling. Dev. Cell.

57. Song, H., Niu, X., Quan, J., Li, Y., Yuan, L., Wang, J., Ma, C., and Ma, E. (2020). Discovery of specific HDAC6 inhibitor with anti-metastatic effects in pancreatic cancer cells through virtual screening and biological evaluation. Bioorganic Chem. 97, 103679.

58. Stefan, E., Aquin, S., Berger, N., Landry, C.R., Nyfeler, B., Bouvier, M., and Michnick, S.W. (2007). Quantification of dynamic protein complexes using Renilla luciferase fragment complementation applied to protein kinase A activities in vivo. Proc. Natl. Acad. Sci. 104, 16916–16921.

59. Streets, A., and Ong, A. (2020). Post-translational modifications of the polycystin proteins. Cell. Signal. 72, 109644.

60. Streets, A.J., Wessely, O., Peters, D.J.M., and Ong, A.C.M. (2013). Hyperphosphorylation of polycystin-2 at a critical residue in disease reveals an essential role for polycystin-1- regulated dephosphorylation. Hum. Mol. Genet. 22, 1924–1939.

61. Sun, L., Hu, C., and Zhang, X. (2019). Histone Deacetylase Inhibitors Reduce Cysts by Activating Autophagy in Polycystic Kidney Disease. Kidney Dis. 5, 163–172.

62. Taskén, K., and Aandahl, E.M. (2004). Localized effects of cAMP mediated by distinct routes of protein kinase A. Physiol. Rev. 84, 137–167.

63. Tunquist, B.J., Hoshi, N., Guire, E.S., Zhang, F., Mullendorff, K., Langeberg, L.K., Raber, J., and Scott, J.D. (2008). Loss of AKAP150 perturbs distinct neuronal processes in mice. Proc. Natl. Acad. Sci. 105, 12557–12562.

64. Wallace, D.P. (2011). Cyclic AMP-mediated cyst expansion. Biochim. Biophys. Acta 1812, 1291–1300.

65. Weber, E.J., Chapron, A., Chapron, B.D., Voellinger, J.L., Lidberg, K.A., Yeung, C.K., Wang, Z., Yamaura, Y., Hailey, D.W., Neumann, T., et al. (2016). Development of a microphysiological model of human kidney proximal tubule function. Kidney Int. 90, 627– 637.

66. Westlake, C.J., Baye, L.M., Nachury, M.V., Wright, K.J., Ervin, K.E., Phu, L., Chalouni, C., Beck, J.S., Kirkpatrick, D.S., Slusarski, D.C., et al. (2011). Primary cilia membrane assembly is initiated by Rab11 and transport protein particle II (TRAPPII) complex-dependent trafficking of Rabin8 to the centrosome. Proc. Natl. Acad. Sci.

67. Wheway, G., Nazlamova, L., and Hancock, J.T. (2018). Signaling through the Primary Cilium. Front. Cell Dev. Biol. 6.

68. Whiting, J.L., Nygren, P.J., Tunquist, B.J., Langeberg, L.K., Seternes, O.-M., and Scott, J.D. (2015). Protein Kinase A Opposes the Phosphorylation-dependent Recruitment of Glycogen Synthase Kinase 3β to A-kinase Anchoring Protein 220. J. Biol. Chem. 290, 19445–19457.

69. Whiting, J.L., Ogier, L., Forbush, K.A., Bucko, P., Gopalan, J., Seternes, O.-M., Langeberg, L.K., and Scott, J.D. (2016). AKAP220 manages apical actin networks that coordinate aquaporin-2 location and renal water reabsorption. Proc. Natl. Acad. Sci. 113, E4328–E4337.

70. Wilson, P.D. (2004). Polycystic kidney disease. N. Engl. J. Med. 350, 151–164.

71. Winyard, P., and Jenkins, D. (2011). Putative roles of cilia in polycystic kidney disease. Biochim. Biophys. Acta BBA - Mol. Basis Dis. 1812, 1256–1262.

72. Ye, H., Wang, X., Constans, M.M., Sussman, C.R., Chebib, F.T., Irazabal, M.V., Young, W.F., Harris, P.C., Kirschner, L.S., and Torres, V.E. (2017). The regulatory 1α subunit of protein kinase A modulates renal cystogenesis. Am. J. Physiol. - Ren. Physiol. 313, F677–F686.

73. Yui, S., Nakamura, T., Sato, T., Nemoto, Y., Mizutani, T., Zheng, X., Ichinose, S., Nagaishi, T., Okamoto, R., Tsuchiya, K., et al. (2012). Functional engraftment of colon epithelium expanded in vitro from a single adult Lgr5 + stem cell. Nat. Med. 18, 618–623.

